# Reward value revealed by auction in rhesus monkeys

**DOI:** 10.1101/2019.12.11.872564

**Authors:** Alaa Al-Mohammad, Wolfram Schultz

**Affiliations:** Department of Physiology, Development and Neuroscience University of Cambridge, Cambridge CB2 3DY United Kingdom

**Author notes:** Corresponding author: email address Wolfram Schultz. Other email addresses* Alaa Al-Mohammad.

**Keywords:** BDM, second-price auction, bidding, ranking, choice The authors declare no conflict of interest.

## Abstract

Economic choice is thought to involve the elicitation of the subjective values of the choice options. Thus far, value estimation in animals has relied upon stochastic choices between multiple options presented in repeated trials and expressed from averages of dozens of trials. However, subjective reward valuations are made moment-to-moment and do not always require alternative options; their consequences are usually felt immediately. Here we describe a Becker-DeGroot-Marschak (BDM) auction-like mechanism that provides more direct and simple valuations with immediate consequences. The BDM encourages agents to truthfully reveal their true subjective value in individual choices (’incentive compatibility’). Monkeys reliably placed well-ranked BDM bids for up to five juice volumes while paying from a water budget. The bids closely approximated the average subjective values estimated with conventional binary choices, thus demonstrating procedural invariance and aligning with the wealth of knowledge acquired with these less direct estimation methods. The feasibility of BDM bidding in monkeys paves the way for an analysis of subjective neuronal value signals in single trials rather than from averages; the feasibility also bridges the gap to the increasingly used BDM method in human neuroeconomics.

**Significance:** The subjective economic value of rewards cannot be measured directly but must be inferred from observable behavior. Until now, the estimation method in animals was rather complex and required comparison between several choice options during repeated choices; thus, such methods did not respect the imminence of the outcome from individual choices. However, human economic research has developed a simple auction-like procedure that can reveal in a direct and immediate manner the true subjective value in individual choices (Becker-DeGroot-Marschak, BDM, mechanism). The current study implemented this mechanism in rhesus monkeys and demonstrates its usefulness for eliciting meaningful value estimates of liquid rewards. The mechanism allows future neurophysiological assessment of subjective reward value signals in single trials of controlled animal tasks.

## Introduction

In 1797, Goethe wanted to sell his epic poem ‘Hermann und Dorothea’, while also revealing how much the publisher truly valued it (Moldovanu and Tietzel 1998). To achieve this, Goethe set a secret reserve price below which he would not sell the poem and asked the publisher for an offer. If the offer was above Goethe’s secret price, Goethe would sell it for that price. The publisher’s optimal strategy was to bid an amount equal to their value; thus, Goethe’s auction achieved the property of truthful value elicitation or ‘incentive compatibility’ (Milgrom and Weber 1982; Karni and Safra 1987). By bidding more than their value, the publisher allowed the possibility of winning the poem at a price greater than their value, thus making a loss. By bidding less than their value, they allowed the possibility of losing the poem even if Goethe’s reserve price was less than their value; and would not reduce the cost, as the cost would always equal the reserve price.

Goethe’s method was a ‘second-price’ auction, whose various forms are staples of experimental economics (Lusk and Shogren, 2007). The Becker-DeGroot-Marschak mechanism (BDM; Becker et al., 1964) is an equivalent ‘incentive compatible’ auction that directly reveals the subjective value of the good for the bidder. In BDM, the bidder’s stated value for a good is compared against a random computer price or bid: the auction is won if the stated bid equals or exceeds the computer bid and is lost otherwise. The BDM is increasingly used in human neuroimaging and allows experimenters to elicit true reward valuations and correlated brain activity (Plassmann et al., 2007; Chib et al., 2009; Linder et al., 2010; Harris et al., 2011; Tang et al., 2014; Tyson-Carr et al., 2018).

Other ‘incentive compatible’ means of value elicitation have depended upon repeated binary choice (BC) between various goods, whereby subjective values of individual goods are inferred from observed choices (Platt and Glimcher, 1999; Padoa-Schioppa and Assad, 2006; Kobayashi and Schultz, 2008; Lak et al., 2014). In BC tasks the chosen option is inferred to have the highest value within the set of competing options or, with indifference, the same value as other options. As fruitful as this approach has been, it may not be without problems. In primates, rational choice is frequently violated (Tversky, 1972; Knetsch and Sinden, 1984; Tversky and Simonson, 1993; Bateman et al., 1997; Rieskamp et al., 2006), neuronal reward signals adapt to current distributions (Tobler et al., 2005; Kobayashi et al., 2010; Louie et al., 2011; Soltani et al., 2012; Padoa-Schioppa and Rustichini, 2014), and choice might involve simple heuristics rather than value comparison (Brandstätter et al., 2006; Vlaev et al., 2011; Piantadosi and Hayden, 2015).

These results suggest that the method of value elicitation may impact on the decision-making process and valuation. It is therefore useful to have multiple value elicitation mechanisms which engage decision-making differently. In perhaps the most striking difference, the agent directly states the subjective value for the good in each BDM trial, while the value in BCs is only inferred from multiple choices. The direct value statement removes potential BC confounds, like complexity of presentation of multiple options, comparison between viable options, and satiety.

Although invasive neurophysiology allows for finer mechanistic resolution, animal experiments have thus far been confined to BC tasks, and a neuronal basis of BDM bidding is missing.

We investigated the feasibility of using the BDM with monkeys. Two rhesus monkeys bid different water volumes for the chance to obtain specific juice volumes in individual BDM trials. They used a joystick to move an on-screen cursor, thus specifying the volume of water they would ‘pay’ for specific juice volumes represented by unique visual stimuli. Both monkeys reliably expressed well-ranked bids, with distinct water values, for each of the five different juice volumes offered. Moreover, subjective values in the BDM were like those inferred from an equivalent BC task.

## Materials and Methods

### Animals and materials

Two purpose-bred and group-housed male rhesus monkeys (Macaca mulatta), A (weighing 10.8kg) and B (weighing 7.9kg), were used for this study. Both animals were trained, via several intermediate tasks, on the BDM and a closely related binary choice (BC) task over a period of 24 and 36 months respectively. The animals participated in experiments for 1-2 hours every weekday.

This research has been approved and supervised by the UK Home Office, UK Animals in Science Committee and UK National Centre for Replacement, Refinement and Reduction of Animal Experiments (NC3Rs), and locally at the University of Cambridge by its Animal Welfare and Ethical Review Body (AWERB), Governance and Strategy Committee, Biomedical Service (UBS) Certificate Holder, Welfare Officer, Named Veterinary Surgeon (NVS), and Named Animal Care and Welfare Officer (NACWO).

### General experimental design

During experimental sessions, monkeys sat in a primate chair (Crist Instruments) positioned 60 cm from a computer monitor. They made choices in the BDM and BC tasks using a custom-built joystick (Biotronix Workshop, University of Cambridge). The joystick allowed for both forward/backward movement to move the bid cursor up/down in the BDM task and left/right movement to choose between the options in the BC task. The joystick also had a touch sensor that detected whether the monkey was holding it.

Joystick position data and digital task event signals were sampled at 2 kHz and stored at 200 Hz (joystick) or 1 kHz (task events). Liquid reward was delivered by a computer-controlled solenoid liquid valve (∼0.006 ml/ms opening time), with a standard deviation of droplet size approximately equal to 0.06 ml. Behavioral tasks were controlled by custom-made software (MATLAB; The MathWorks) running in conjunction with the Psychophysics toolbox (Brainard, 1997) on a Microsoft Windows 7 computer.

### Becker-DeGroot-Marschak (BDM) task design

On each BDM trial, the monkey competed with a randomly set computer bid for obtaining a specific volume of fruit juice (the same 10% apple and mango concentrate was used as the juice reward in all tasks). If the monkey’s bid equalled or exceeded the computer bid, it received the specified fruit juice volume and paid the price indicated by the computer bid from the water budget that reset in each trial to 1.2 ml. If the monkey’s bid was lower than the computer bid, it did not receive the fruit juice volume but kept the water budget that was delivered at trial end.

Monkeys bid with a forward-backward moveable joystick for specific volumes of fruit juice. The procedure involved five fractals, each indicating one of five specific volumes of the same fruit juice (Fig. 1*A*). Only one fractal was shown on any given trial. A grey rectangle represented the water budget (Fig. 1*B*). A red bar on the grey rectangle indicated the joystick position, and thus the monkey’s bid, on a computer monitor in front of the monkey, and a green bar on the same grey rectangle indicated the computer bid (higher was more). The left side of the computer monitor displayed the fractal, and the right side displayed a grey rectangular ‘budget bar’ whose total area represented the 1.2 ml of water with which the monkey could bid on each trial (Fig. 1*C*). The monkey placed a bid by moving the red cursor up/down by pushing/pulling the joystick. Following the monkey’s bidding period, the computer-bid appeared (green line). If winning the BDM (Fig. 1*C*, top), a volume of water equivalent to that remaining above the green line was delivered first, followed later by the juice volume indicated by the fractal; thus, the water volume occluded below the computer’s bid was the price paid for the juice gained. If losing the BDM (Fig. 1*C*, bottom), then the fractal representing the juice reward disappeared and the full 1.2 ml water budget was delivered, but this was not followed by any juice.

**Figure 1.**
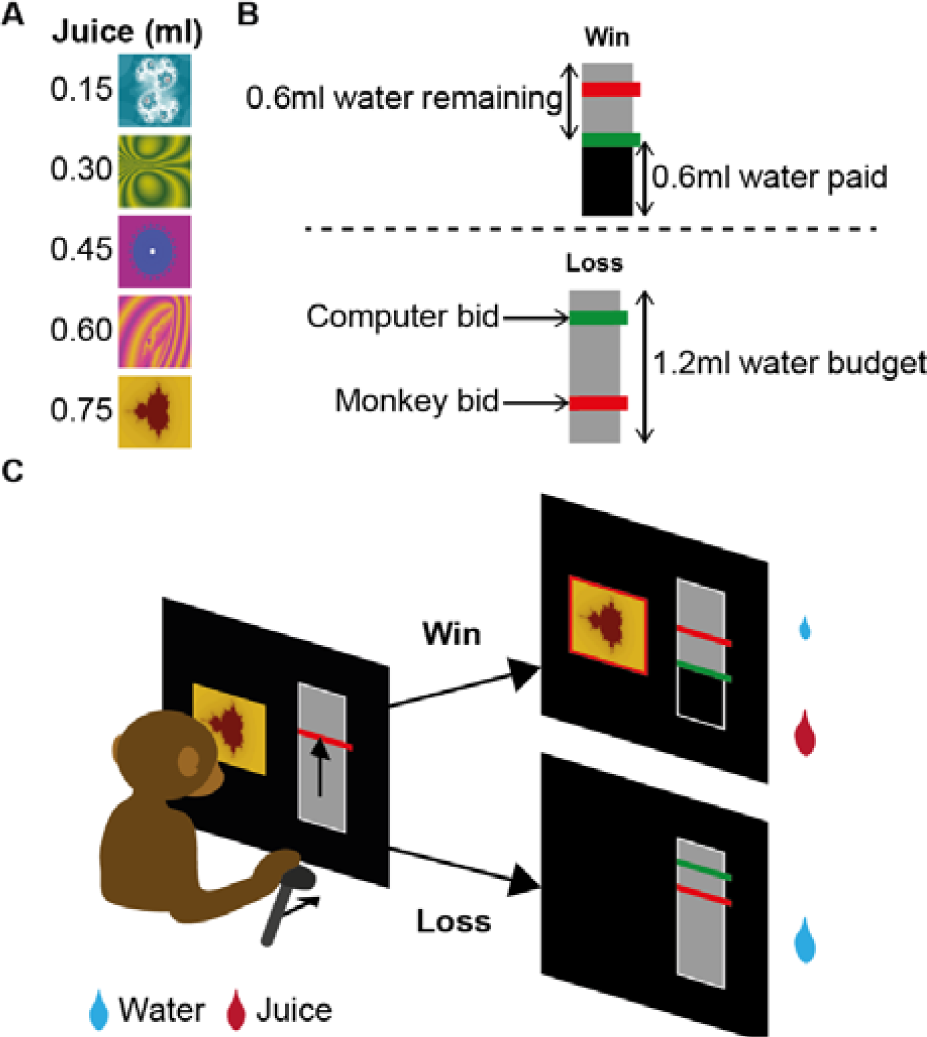
A BDM task for monkeys. *A*, Five fractals indicating five specific volumes of same fruit juice. ***B,*** Gey rectangles indicating pay-out of water budget. Each trial started with a new water budget of 1.2 ml. Monkey bids and computer bids are indicated by red and green bars, respectively (higher = more). Top: winning the auction (monkey bid ≥ computer bid) resulted in receiving the water volume remaining above the green bar (second price auction) plus the desired fruit juice volume 0.5 s later. Bottom: losing the auction (monkey bid < computer bid) resulted in receiving the full water budget but no fruit juice. ***C*,** Bidding task. The monkey placed a bid by moving the red cursor up-down via pushing-pulling a joystick. Then the computer bid appeared (green bar). Then the BDM result was paid out as described in (B).

Trials were independent of one another, and each monkey completed 200 correct BDM trials in each daily session of testing. Because the monkeys’ bidding behavior might be explained by motor vigor or simple conditioned motor responses, we trained them using three different starting positions for their bid cursor; either at the bottom (B), top (T), or, at a random position (R) on the budget bar. Monkeys completed 10 sessions under each of these 3 conditions, for a total of 30 BDM sessions for each monkey.

### BDM trial structure

The beginning of each BDM trial was signalled to the monkey by a yellow cross at the center of the monitor during a 0.5 s Preparation epoch (Fig. 2*A*). This was followed by an Offer epoch with presentation of the juice volume to bid for, represented by a specific fractal image, and the rectangular ‘budget bar’ stimulus whose total grey area indicated 1.2 ml of water. A dark-red horizontal bar (bid cursor) also appeared within the confines of the budget bar. The Offer epoch was presented for a variable time, mean 2 s ± 1 s with a flat hazard rate, as such temporal uncertainty is known to encourage attention to stimulus changes.

**Figure 2.**
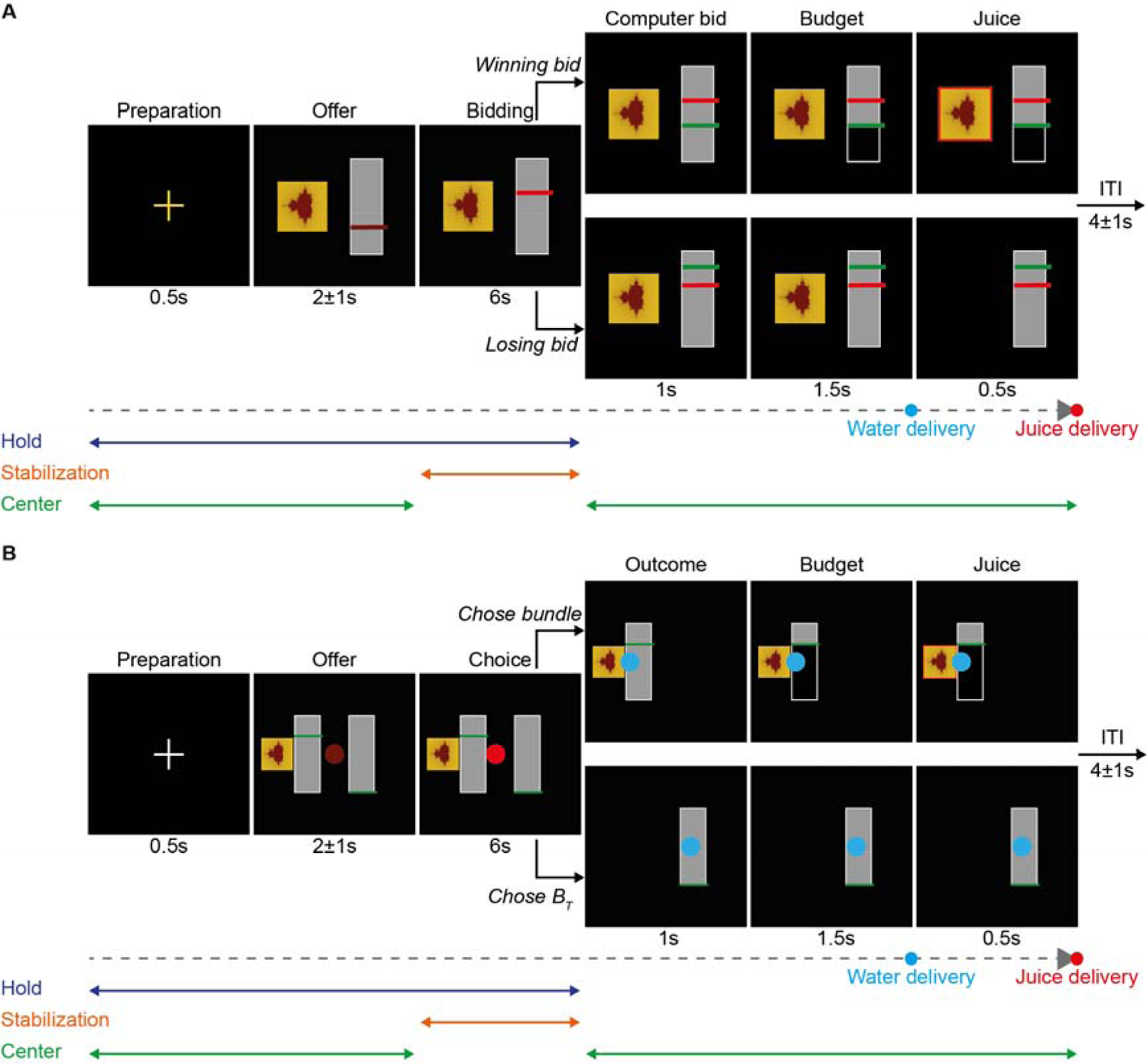
Timeline of BDM and Binary Choice (BC) tasks. *A*, BDM task. A cross during the Preparation epoch prompts the monkey had to maintain grasp of a joystick (blue line, ‘Hold’) and keep it in a central position (left green line, ‘Center’). In the subsequent Offer epoch, the monkey was presented with a fractal image indicating the volume of juice to bid for; the full water budget; and the bid cursor’s starting position. The Bidding epoch began after a variable delay governed by a flat hazard function. Now the monkey was free to move the red bidding cursor via the joystick within the grey vertical rectangle. Each bid was made by the monkey stabilizing the cursor at the desired position for >250 ms after it had moved it there to place a bid (orange line, ‘Stabilization’). Failure to make a bid within the 6s Bidding period, or joystick release before the end of this period, resulted in trial termination and constituted an error. Joystick movement outside the Bidding epoch also constituted an error. The computer bid was displayed after the Bidding epoch (and the monkey turned the joystick-cursor back to the central position and held it there without moving the cursor, right green line, ‘Center’). If the monkey’s bid was higher than the computer’s (win), the budget bar below the computer bid was occluded and the monkey received the remaining water budget at the end of the Budget epoch, and the juice at the end of the Juice epoch. Otherwise (loss), the full 1.2 ml water budget was delivered at the end of the Budget epoch, but no juice was delivered. Trials were separated by a variable inter-trial interval (ITI) of 4 ± 1 s. ***B*,** BC control task. Stimuli, rewards, delays after stimuli and movements were the same as in the BDM. The same behavioral requirements applied at equivalent epochs (blue, orange and green lines): centring of joystick in the Offer epoch; stabilising of bid cursor position in the Bidding epoch; and no joystick movement allowed outside of the Bidding epoch.

After the Offer epoch, monkeys used the joystick to move the bid cursor up/down within the confines of the budget bar. The beginning of this Bidding epoch was indicated by a color change of the bid cursor. Monkeys had 6s to place a bid and did so by maintaining a given bid cursor position for >0.25 s. Following stabilization of the bid cursor’s position, it could no longer be moved. The monkey waited until the end of the 6s bidding period regardless of when it had finalized its bid.

Thus, the monkey could not manipulate reward rate or temporal reward discounting by making bids more/less quickly. Failure to stabilize their bid cursor within the 6s Bidding epoch resulted in abortion of the trial. Bidding was followed by a Computer Bid epoch in which a green horizontal bar (computer bid cursor) appeared within the budget bar at a position corresponding to the randomly generated computer-bid for that trial. Trials were interleaved with intertrial intervals of random duration (4 s ± 1 s, conforming to a truncated exponential function).

### Computer bid distribution

Following extensive modeling, we selected a beta distribution of computer bids over a uniform (flat) distribution to encourage more accurate bidding. A beta distribution does not compromise the crucial incentive compatibility of the BDM (obtaining optimal outcome by truthfully bidding according to own value) but incurs a higher cost of inaccurate bidding (’cost of misbehavior’) compared to a uniform distribution without changing the points of maximal expected and observed payoff (see below Fig. 9 and Extended Data Fig. 9-1) (Lusk and Shogren, 2007; Tymula et al., 2016). Thus, we generated computer bids from a pseudo-normal beta distribution, with support [0,1] and parameters (α = 4, β = 4), mean = 0.5 and variance = 0.028; the random number thus generated was simply multiplied by the maximum bid of 1.2 ml to generate a bid between 0 ml and 1.2 ml. Presentation of the computer bid was followed by a 1.5 s Budget epoch: if the monkey’s bid was higher than the computer’s, then the water budget to be paid was represented by occluding the area between the bottom of the budget bar and the computer’s bid cursor; otherwise, there was no change in the display as no payment was required. In either case the remaining volume of water was delivered at the end of the Budget epoch. Finally, trials ended with a 0.5 s Juice epoch which followed the onset of water delivery by 0.5 s. If the monkey had made a winning bid, then the fractal was surrounded by a red border and the indicated volume of juice was delivered. Otherwise, the fractal disappeared, and no juice was delivered at the end of the Juice epoch.

For a given bid on an individual trial, the probability of winning the BDM, as well as the expected remaining water budget, are determined by the distribution from which the computer bid has been drawn. Thus, BDM bidding involves uncertainty. The bidder may learn the probability distribution of winning and losing the BDM in each trial; this aspect can be described as being essentially risky. With on-going experience, the uncertainty of computer bids changes gradually from initial ambiguity (when the distribution is only incompletely known) towards risk (when the distribution of computer bids becomes completely known), although it may reasonably be doubted that an animal can learn the distribution completely given the various demanding factors of the BDM task. Thus, an element of ambiguity (true uncertainty) may always remain present in animals performing in a BDM task. Such elements of ambiguity and risk are not present in equivalent BC tasks (unless they have been specifically designed so). This characteristic may make BDM performance cognitively more demanding and, depending on ambiguity and risk attitude, potentially less desirable than BC tasks (Platt and Huettel, 2008).

### Error trials

Monkeys were required to maintain hold of the joystick from the Preparation epoch to the end of the Bidding epoch, and to always maintain the joystick in a central position, except during the Bidding epoch. Failure to comply with these restrictions was considered an error and led to abortion of the trial. All errors resulted in the same blue error monitor, error sound, and a delay of 3 s plus the remaining trial time with no further reward delivery.

Across the 30 sessions of BDM testing, Monkey A made 433 errors out of 6,433 trials (6.73%), and Monkey B made 2,692 errors out of 8,692 trials (30.97%) (Extended Data Tables 2-1 and 2-2). However, most of Monkey B’s errors consisted of long strings of consecutive trials during which the animal failed to attend to the task, which may have been facilitated by the lack of some constraints: the animals were free to move their head and /gaze away from the monitor.

Specifically, Monkey A made an average of 14.43 ± 20.04 errors per session (range 0 - 97), giving a mean error rate of 6.08 ± 7.23% across all sessions (range 0 - 23.66%). In the B-BDM, monkey A’s errors were predominantly due to not maintaining hold of the joystick and made up 55.09 ± 36.11% of all errors in these trials, while these accounted for only 2.95 ± 7.88% of errors in the T-BDM and 12 ± 17.79% of errors in the R-BDM. Overall, Monkey A’s error rate improved over the course of the sessions. In the first 10 sessions (B-BDM), Monkey A’s mean overall error rate was 12.88 ± 8.89% but over the last 10 sessions (R-BDM) the overall error rate was 1.65 ± 1.68%. Accordingly, for Monkey A there was a significant negative Pearson’s correlation between the number of errors made in a session and the session number: Rho = −0.60, p = 4.25×10^-4^.

Monkey B performed significantly worse than monkey A and did not appear to make any such improvement over the course of the 30 sessions; there was no correlation between the number of errors made in a session and the session number for Monkey B, with Pearson’s Rho = −0.066, p = 0.73. Overall, monkey B made a mean average of 89.73 ± 49.42 errors per session (range 17 - 199), giving a mean error rate of 29.09 ± 11.61% across all sessions (range 7.83 - 49.87%).

For Monkey A, errors tended to occur on individual trials, with few trials with consecutive errors; the average length of consecutive errors for monkey A (i.e. including only errors that were followed by another error) was 1.26 ± 1.25 across all sessions, and this animal never had more than 5 consecutive error trials. By contrast, Monkey B at times failed to attend to the task for long strings of consecutive trials; reflecting this, the average length of consecutive errors for Monkey B was 3.11 ± 1.04, with up to 30 consecutive error trials in one session (session 23).

Analysis of joystick movement also suggested that most errors were due to inattention to the task for both monkeys. For Monkey A, there was no movement of the joystick in 400 of 433 error trials (92.4% of error trials), and for Monkey B there was no movement of the joystick in 2266 of 2655 error trials (85.35% of error trials).

### Binary Choice (BC) procedure

The most important factor motivating the design of our stochastic BC task was the elicitation of value for comparison with BDM bids while maintaining a perceptual and economic equivalence between the tasks. Thus, the same stimuli and rewards were used in both tasks, and the timings of analogous stimulus changes, choice periods, behavioral requirements, and reward events were the same between them (Fig. 2*B*).

The beginning of each BC trial was signalled by a white cross at the center of the monitor during a 0.5 s Preparation epoch. This was followed by an Offer epoch with presentation of two options on either side of the monitor: one of the options consisted of a bundle formed of a specific juice volume (indicated by a specific fractal) together with a variable volume of water budget (quantitatively indicated by the grey area above the green line), and the other option consisted of the fixed full water budget (B_T_; indicated by the full grey rectangle). The side on which each of these options appeared was randomized on each trial. A dark-red circle (choice cursor) also appeared at the center of the monitor. The Offer epoch was presented for a variable time, with mean 2 s ± 1 s with a flat hazard rate.

After the Offer epoch, the monkey used the joystick to move the choice cursor left/right within the confines of the monitor. The beginning of this Choice epoch was indicated by a color change of the choice cursor. The monkey had 6s to make a choice and did so by maintaining a given choice cursor position for >0.25 s, choices also had to fall within the rightmost/leftmost third of the monitor, where the choice cursor changed color from red to blue. Following stabilization of the choice cursor’s position, it could no longer be moved. The monkey had to wait until the end of the 6s choice period regardless of when they had stabilized the choice cursor, and so could not alter reward rate or temporal reward discounting by making choices more/less quickly. Failure to stabilize their choice cursor within the 6 s Choice epoch resulted in abortion of the trial with an error.

The Choice epoch was followed by a 1 s Outcome epoch, which began with the unchosen option disappearing from the monitor. After this, the 1.5 s Budget epoch began: if the bundle was chosen then the water budget difference between the bundle and B_T_ was occluded at the beginning of this epoch, otherwise, if the monkey had chosen B_T_, then no further stimulus changes took place.

In either case the volume of water indicated by the chosen option was delivered at the end of the Budget epoch.

Finally, trials ended with a 0.5 s Juice epoch which immediately followed water delivery. If the monkey had chosen the bundle, then the fractal was surrounded by a red border and the indicated volume of juice was delivered. Otherwise, no stimulus change took place, and no juice was delivered at the end of the Juice epoch.

Trials were interleaved with inter-trial intervals of random duration (4 s ± 1 s, conforming to a truncated exponential function). The monkeys were required to maintain hold of the joystick from the Preparation epoch to the end of the Choice epoch, and always had to maintain the joystick in a central position, except during the Choice epoch; all other behaviors were considered as errors and led to the trial being aborted. All errors resulted in the same blue error monitor, error sound, and a delay of 3 s plus the remaining trial time without further liquid delivery.

Monkey A made 378 errors in 2,378 BC trials (15.90%) and Monkey B made 721 errors in 2,721 trials (26.50%). For both monkeys most errors were due to long strings of consecutive trials during which they did not engage in the task.

### Stimulus training

We trained each monkey to associate fractal visual cues with different volumes of the same juice over a period of 2 months of daily training (Fig. 3*A*). At this stage, the monkeys were also trained to maintain hold of the joystick for each trial to progress to juice delivery. This hold requirement was used in all subsequent training procedures and both the BDM and BC tasks as detailed above.

**Figure 3.**
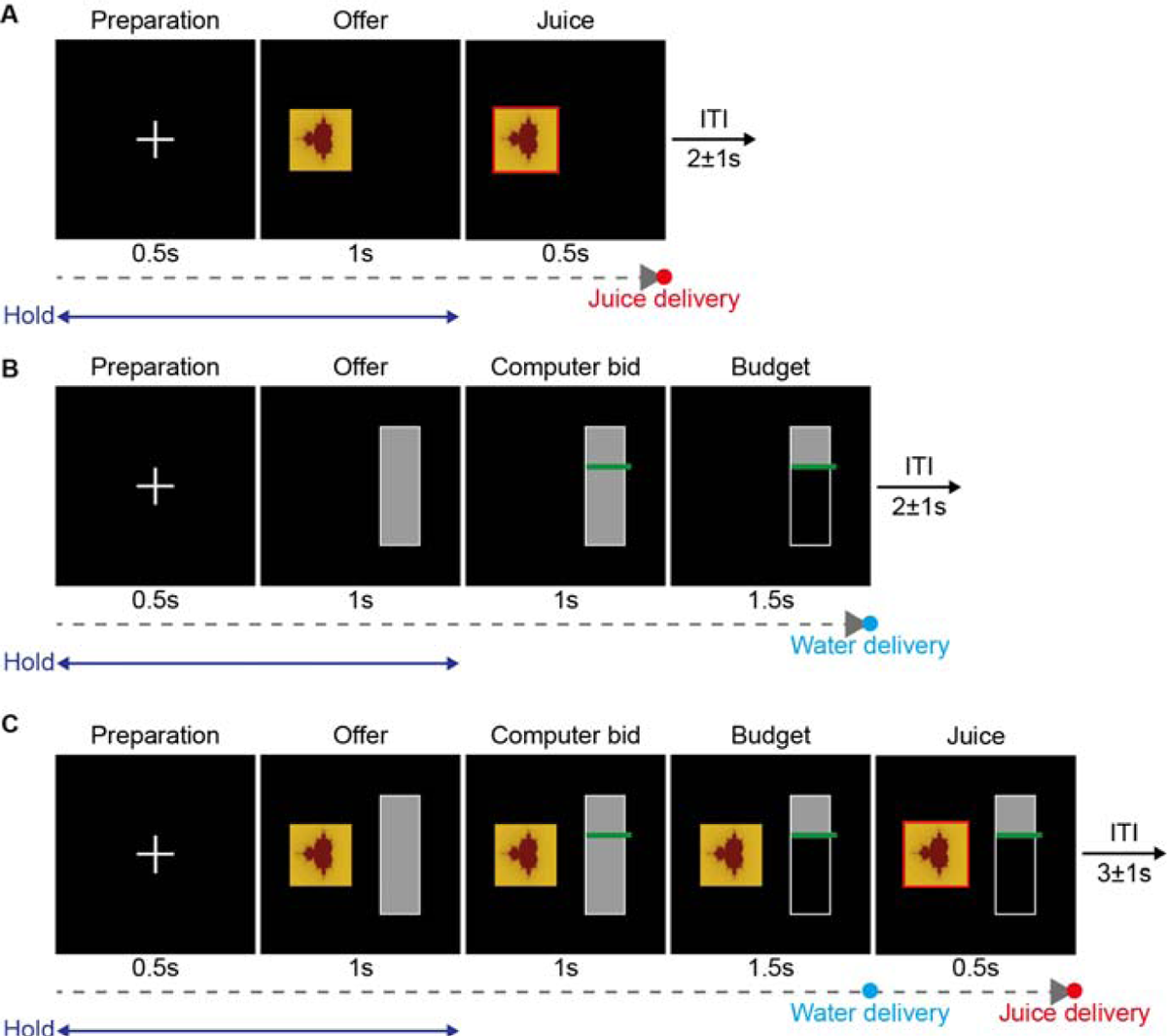
Stepwise learning of stimulus-juice associations. *A*, Initial learning to associate each of 5 unique fractal images with 5 specific juice volumes. Fractals were surrounded by a red border 0.5 s before juice delivery, as in the final BDM and BC tasks. At this point, the monkey was also taught to maintain hold of the joystick throughout Preparation and Offer epochs (blue line, ‘Hold’); else trials were considered erroneous and aborted. Monkeys each completed 20 of these training sessions to learn the stimuli associated with each of the 5 juice volumes used. ***B*,** Subsequent learning to associate the budget bar with water budget volumes. The monkey was presented with a grey bar stimulus whose full area represented 1.2 ml of water. Then a green cursor, as later used to indicate the computer bid in the BDM, appeared at a random location on the vertical rectangle, and the area of the rectangle below was occluded. The monkeys received the remaining volume of water (% of remaining grey area × 1.2 ml) at 1.5 s after occlusion of the rectangle below the computer bid cursor, as in the final BDM and BC tasks. Monkeys each completed 20 of these training sessions to learn the budget bar stimulus contingencies. ***C*,** Learning the relative timing of delivery of water budget and juice. The monkey was presented with both stimuli concurrently. Both the BDM and BC tasks had identical timing of water delivery (from the point at which the budget bar was occluded below the green cursor) and juice delivery (0.5 s later). Monkeys each completed 20 sessions of these training sessions.

The monkeys then learnt to associate the grey area of the budget bar with a corresponding *v*olume of water over another month of training (Fig. 3*B*). On each trial, the green cursor stimulus used to indicate computer bids in the BDM task appeared at a random location on the budget bar, and the area of the bar below this was occluded. The monkeys received a volume of water proportional to the remaining grey budget area, with the full area predicting 1.2 ml of water.

We then trained the monkeys in sessions in which both the juice and water budget appeared concurrently over a period of approximately 1 month (Fig. 3*C*). The indicated volumes of water and juice were then delivered in the same order and with the same delay that would be used in the BDM task.

### Joystick training

After the monkeys had learnt the stimulus-reward associations, they were trained to operate the joystick over a period of 3 months. For left/right movement, monkeys were first trained on a binary choice task, with budget bars presented on either side of the monitor (Fig. 4*A*). On each trial, monkeys had to move a red circular cursor from the center of the monitor to their preferred side within a 6s choice epoch. The cursor changed color from red to blue at the rightmost or leftmost third of the monitor to indicate that the cursor had been moved far enough to choose the offer on that side. The monkeys then had to stabilize the cursor in a given position to indicate that a choice had been made, else the trial would end with an error. We started by presenting budget bars offering large differences in water volume and gradually reduced the difference in volume between the two offers as the animals came to reliably choose the option with the most water. This task taught the monkeys left/right movement of the joystick for the final BC task and confirmed that they understood the significance of the grey area of the budget bar. We also trained the monkeys in a version of this task in which both stimuli offered the same volumes of water, such that neither should be preferred. Given the lengthy periods of joystick training, we had time to eliminate any side bias, and we altered the joystick gain in left/right directions such that each of the left and right equally rewarded options were chosen with equal probability.

**Figure 4.**
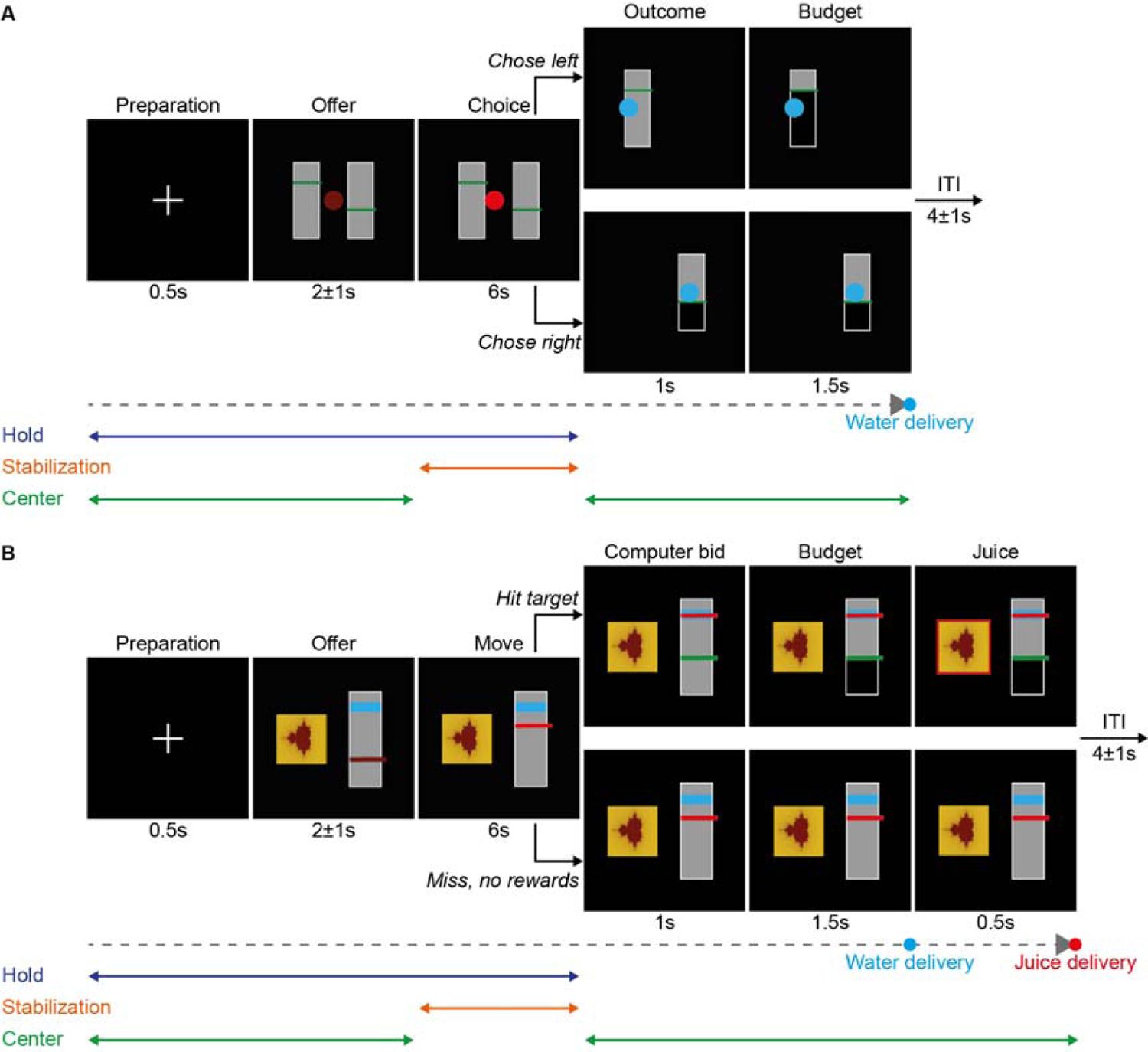
Joystick training tasks. *A*, Horizontal movement. Monkeys moved a red circle on the monitor by making left/right movements of the joystick from a central holding position. The monkey moved the circle into the left or right third of the monitor and stabilised its location for 250 ms, by recentering the joystick, to state its choice. Each monkey performed this task with two different water volumes on either side, as such this task also confirmed that monkeys’ understanding of the budget bar stimulus. On a subset of these trials, we eliminated any possible choice bias by adjusting the gain of joystick movement on either side until identical water volumes were chosen with equal probability. Monkeys each completed 20 of these training sessions to learn left/right movement of the joystick for performance of the BC task. Monkeys were considered to have learnt the task to a sufficient degree when they chose the strictly better option >95% of the time in each direction. **B,** Vertical movement. With similar task epochs and task requirements as the initial choice task (blue, orange and green lines). The monkey was taught to move a cursor vertically on the monitor with forward/backward movements of the joystick. The monkey had to move a red cursor into a randomly positioned blue target area. If it placed the cursor successfully into the target area, the computer bid appeared, and the monkey received the juice and water after the same delay and with the same contingencies as in the BDM task (i.e. only receiving the juice and paying losing some water if their bid was greater than the computer’s bid). If the cursor was not secured within the target area in the Move epoch, then no further stimulus change took place until trial end, and all rewards were withheld. The height of the blue target area was progressively reduced as the monkey’s performance improved, until the target area was 1/10^th^ of the height of the budget bar. Note that these training sessions were essentially equivalent to the final BDM task but with forced bids. Monkeys each completed 40 of these training sessions to learn accurate up/down movement of the joystick for use in the BDM task. Monkeys were considered to have learnt the task to a sufficient degree when they could accurately place their bid in the target region >95% of the time for each location in the budget bar (targets appeared across the budget bar in each of 10 equally sized target areas).

Finally, the monkeys were trained to move the bid cursor by moving the joystick forwards/backwards. They performed a task in which there were both juice and budget bar cues, like the final BDM task, however, in this case they had 6s to move the bid-cursor into a blue target area which appeared at a random location on the budget bar (Fig. 4*B*). The bid cursor had to be stabilized within the target area, else the trial would end due to failure to meet the stabilization requirement. This would therefore act as a forced bid, and the rest of the trial proceeded as in the BDM task, with the appearance of a green cursor at a random height and receipt of either some water and juice or the full volume of water, depending on the relative locations of the monkey’s red cursor and the randomly generated green cursor. As monkeys’ performance improved, we gradually decreased the size of the blue target’s height, until they could reliably perform the task with a target that was 1/10^th^ of the total budget bar height.

### Joystick control

Voltage outputs for joystick movement in both axes were separate, and in the central position the voltage output was 0 V. A maximal forward or rightward movement produced an output of 5 V, and a maximal backward or leftward movement produced an output of −5 V. The positions of on-monitor cursors were modulated by the following equations, where ***G*** is the gain or amplification applied to the voltage modulation, ***V***, and ***P*** is the pixel position of the center of the cursor at time ***T***:

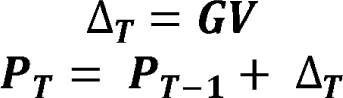

Thus, the value of ***P*** changes more quickly with greater deflections of the joystick. In the BDM, forward and backward deflections of the joystick move the bid cursor up and down the budget bar, with the maximum and minimum values of ***P*** being limited to the top and bottom pixel positions of the budget bar. In the BDM, the value of ***G*** was the same for movements in both directions.

In the BC task, the value of ***G*** depended on whether ***V*** took a positive or negative value, thus the gain could be set differently for rightward/leftward joystick movements. This feature counteracted the effects of side-bias on the animal’s choices. Values of ***G*** were set for each direction such that the monkeys made choices without a statistically significant side-bias when both the left and right-hand-side offers were the same.

The monkeys found it difficult to hold the joystick perfectly still in the central position, so a window of tolerance for slight movements was necessary to prevent small erratic deflections of on-monitor cursors during choice/bidding epochs. A minimum threshold of 2% of the maximal voltage displacement was applied in every direction, such that any output with an absolute magnitude of 0.1 V or less was treated as a 0 V modulation and did not produce any deflection of on-monitor cursors.

For tight control of monkeys’ movements, we enforced three behavioral requirements relating to joystick control, failure of which led to a blue error monitor for a duration equal to the remaining trial time plus 3s, and with no reward for that trial:

– Hold requirement: The monkeys had to maintain hold of the joystick throughout choice/bidding epochs and in all epochs preceding them, as detected by a built-in touch sensor.
– Center requirement: The monkeys had to maintain the joystick in a central position outside of the choice/bidding epochs, such that only deflections leading to voltage outputs less than or equal to 0.1 V were tolerated in all other epochs.
– Stabilization requirement: The monkeys had to stabilize on-monitor bid and choice cursors in their desired final position for 250 ms, such that the voltage output was less than or equal to 0.1 V for 500 consecutive samples at 2 kHz. This indicated a purposeful choice and had to be completed within the 6s allocated to the choice/bidding epochs.

### Statistical Analysis of BDM bids

To evaluate how well monkeys’ bids reflected increasing juice volumes on individual days of BDM testing, or ‘sessions’, we used Spearman rank correlation (MATLAB: corr) between bids and juice volumes that assumes a monotonic, but not necessarily linear, relationship between the two variables. The bids in BDM reflect the subjective reward value of the item the agent bids for or, more formally, the economic utility of the item. However, the metric of utility in our BDM was the physical amount (milliliters) of the water the animal ‘paid’ from the budget, rather than being utils. To express subjective value properly in utils would have required estimation of a utility function for water separately for each animal. While our monkeys show different degrees of nonlinearity in riskless and risky utility functions for various liquids (Stauffer et al., 2014; Genest et al., 2016; Bujold et al., 2021), the current absence of an empirically estimated utility function for water would render the utility of the measured bids basically uninformative. Nevertheless, we aimed to obtain supplementary confirmatory information on the bid-juice relationship by using other models, including a linear function, two power functions and a 2-parameter Prelec function (Prelec, 1998). As we pooled across different sessions for these preliminary analyses, we used mixed-effects models with a fixed-effect of reward volume and a random-effect of session number (we performed further analyses on individual sessions, as values and preferences may vary from between days of testing). We tested both linear mixed-effects models (MATLAB: fitlme) and non-linear mixed-effects models (MATLAB: fitnlm).

We also wanted to assess how distinct monkeys’ mean bids were for different juice volumes in individual sessions. We used 1-way ANOVAs (MATLAB: anova1) to test whether mean bids for different juice volumes were different to one another in each of the 30 BDM sessions. For these and all other ANOVAs, we also present the omega-squared (ω^2^) measure of effect size for different factors. Post-hoc Bonferroni tests for multiple pairwise parisons (MATLAB: multcompare) were performed to find which juice volumes received mean bids that were significantly different to one another, thus reflecting how well monkeys’ bids discriminated different juice volumes.

Within those sessions in which monkeys’ mean bids reliably discriminated all five juice volumes (i.e. all sessions for Monkey A and 21/30 sessions for Monkey B), we identified how quickly they achieved this. We found the first trial, *T_n_*, for which a 1-way ANOVA and Bonferroni-corrected multiple comparisons tests over mean bids were significantly different for all juice volumes and had also been significant for the 10 trials preceding this, *T_n-10_* - *T_n_*.

We performed an unbalanced two-way ANOVA (MATLAB: anovan) on monkeys’ bids with main factors of juice volume and bid starting position condition to explore the relative influence of motor contingencies, which vary with starting position. To interrogate the effects of the starting location of the bid cursor on monkeys’ final bids more closely, we performed a multiple regression analysis (MATLAB: fitlm) on bids, with regressors for the juice volume (JV) and the interaction between each juice volume and the bid cursor’s exact starting position (SP_JV=Xml_), according to Eq. 1. For each animal, this regression analysis was conducted separately for each of the 10 random starting position sessions, finding the mean value of the coefficient for each regressor across sessions. As bid cursor position was expressed in terms of the corresponding bid volume, all regressors had the same units and scale and could therefore be compared directly (see Results). For Monkey A, B_0_ = 0.05 ± 0.1 (mean ± standard deviation, SD); B_1_ = 1.38 ± 0.14; B_2_ = − 0.11 ± 0.12; B_3_ = −0.17 ± 0.1; B_4_ = −0.04 ± 0.06; B_5_ = 0.02 ± 0.05; B_6_ = −0.02 ± 0.04. For Monkey B, B_0_ = −0.03 ± 0.07; B_1_ = 1.42 ± 0.24; B_2_ = 0.04 ± 0.07; B_3_ = −0.02 ± 0.05; B_4_ = 0 ± 0.05; B_5_ = 0.02 ± 0.1; B_6_ = 0 ± 0.16.

### Statistical Analysis of BC value estimation

We used choices the BC task to estimate the water equivalents of different apple and mango juice volumes. Using a logistic regression model, we estimated regression by fitting the probability of choosing the full 1.2 ml water budget, *P (B choice)*, for each of the bundles, which contained variable water volumes, *B_x_*. Each bundle in this analysis was expressed in terms of the difference in water volume between it and the full budget option, Δ*B = B - B_x_*.

For each of the 5 volumes of juice, we fitted the logistic function (MATLAB: fitglm) of the following form onto the choice data from the BC task:

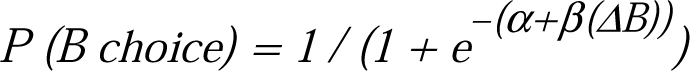

The value of Δ*B* at which *P (B choice)* is equal to 0.5 is an estimate of the monkey’s water-value for the volume of juice which appeared in that set of bundles. In this case, *α* is a measure of choice bias and *β* is a measure of the monkey’s sensitivity to changes in the volume of water available in the budget options. Note, even if Δ*B* is replaced by the ratio of water volumes in the bundle and full budget option, as is the case in some binary choice analyses, we arrive at the same estimates of water value because the volume of water in the budget-only option is constant in this task.

We conducted this analysis on each of the 10 BC sessions for each monkey (Fig. 8*A, B*), but choices were too variable and trials too few to attain reliable value estimates using individual sessions. The monkeys were tested in five BC sessions preceding BDM testing and five BC sessions after BDM testing to detect any change in the values of the juice volumes across the period of BDM testing. No significant change in mean value estimates was detected. We therefore pooled all 10 BC sessions for each animal to acquire better estimates of their average values for these five juice volumes, using the method described above. These acted as our best estimates of the monkeys’ values.

If BC value estimates are taken as the monkeys’ true values for each juice volume, then the optimal bid should be equal to the BC value estimate, except where the estimated value is greater than the maximum bid of 1.2 ml, in which case the optimal bid is equal to this maximal volume. This was only the case for Monkey A’s value for the 0.75 ml apple and mango juice.

How well monkeys’ bids reflected the BC value estimates was determined using a simple linear regression (MATLAB: fitlm) on bids with the BC value estimates for each juice volume as the sole predictor (see results).

The BC value estimates were also used to compute each monkey’s total payoff in terms of water for each trial, as well as the payoffs of optimal and random simulated bidders (see results and following section on simulation methods). This was not possible for the 0.75 ml juice volume, for which Monkey A’s value could not be identified, and as such trials for that juice volume were excluded from those analyses.

### Simulated Bidding

We simulated four types of decision-maker for the BDM task: an optimal decision-maker who always bid the monkey’s exact BC value for each juice volume; a random decision-maker who always made a completely random bid drawn from a uniform distribution with support [0, 1.2]; an over-bidding decision-maker who always bid the monkey’s exact BC value plus 0.2 ml; and an under-bidding decision-maker who always bid the monkey’s exact BC value minus 0.2 ml.

These simulated bidders were presented with the same juice presentations that each monkey faced over 30 BDM sessions of 200 trials each (though trials in which the 0.75 ml juice was presented were excluded for Monkey A as his value for that juice volume and therefore the payoffs, could not be computed - see above). The computer bids for each juice volume were the same as those that each animal faced. BC values were substituted for juice volumes so that payoffs were always expressed in terms of the equivalent volume of water. The mean per-trial payoff was then calculated for each juice volume by dividing the total payoff for that reward by the number of times that reward was presented. This process was repeated separately for each animal.

These simple simulations provided an idea of how each monkey performed in terms of behaviorally relevant outcomes, on a spectrum from completely random behavior to mechanically perfect rational bidding (i.e. with no motor or decision noise), taking the BC values as our best independent estimate of the monkey’s water value for each juice reward.

### Juice-delivery error

To deliver juice and water in our tasks we used a solenoid delivery system, with opening time controlled by voltage pulses. There was an approximately linear relationship between solenoid opening time and the volume of water/juice delivered, and we tested and calibrated the opening times so that we could deliver the appropriate volumes of the different liquids in the task. Calibration of the solenoid systems showed a mean standard deviation of 0.06 ml at any given opening time.

This degree of variability in the volume of liquid delivered at a given solenoid opening time could limit the animal’s ability to discriminate the small differences in expected payoffs that result from different bids in the BDM, as these variations in liquid volume may be indistinguishable from the variability of the solenoid itself.

Increasing water budget volume and juice volume reduces the relative magnitude of the solenoid’s variability in liquid delivery, as the standard deviation of the delivered volume is the same regardless of the mean volume delivered.

These considerations motivated the use of larger liquid volumes in the BDM task. With a larger water budget volume, expected losses are greater for the same pixel distance displacement of the bid cursor from the optimal bid, and the relative contribution of variability in the solenoid delivery is reduced. Thus, monkeys should be able to discriminate differences in expected payoff at smaller relative distances between the actual and optimal bids.

## Results

### Analysis of BDM bids

#### Rank ordered bidding

Once BDM training was concluded, we advanced to testing performance in the BDM task using the final version of the task design detailed above. Fig. 5 shows mean bids from all sessions in both monkeys and post-hoc comparisons of means. For both animals, there were significant differences between bids for the five juice volumes with all three starting positions of the bid cursor (bottom: B-BDM; top: T-BDM, random: R-BDM; one-way ANOVA in each of the 30 sessions, *P* < 0.05: Monkey A: F = 176.42 to 392.36; Monkey B: F = 40.17 to 166.76; Extended Data Table 5-1). Post-hoc t-tests (Bonferroni-corrected for multiple comparisons) confirmed significant differences in all pairwise comparisons of mean bids for the five juice volumes in each of the 30 BDM sessions for Monkey A (all *P* < 0.05), and in 21 of the 30 sessions for Monkey B (*P* < 0.05). With Monkey B, bids differed significantly with all but one pair of juice volumes in eight sessions and two pairs in one session. There were no specific factors explaining Monkey B’s low, although still mostly differential, bids in the early sessions with the top cursor starting position (Fig. 5*B*, T-BDM, sessions 11 - 14). Possibly the low bidding was carried over from the preceding sessions with cursor starting positions at the bottom (B-BDM); the bids recovered subsequently with increasing experience from the more frequent BDM loss from under-bidding (sessions 15 - 20). Taken together, the monkeys made distinct but noisy bids for different rewards.

**Figure 5.**
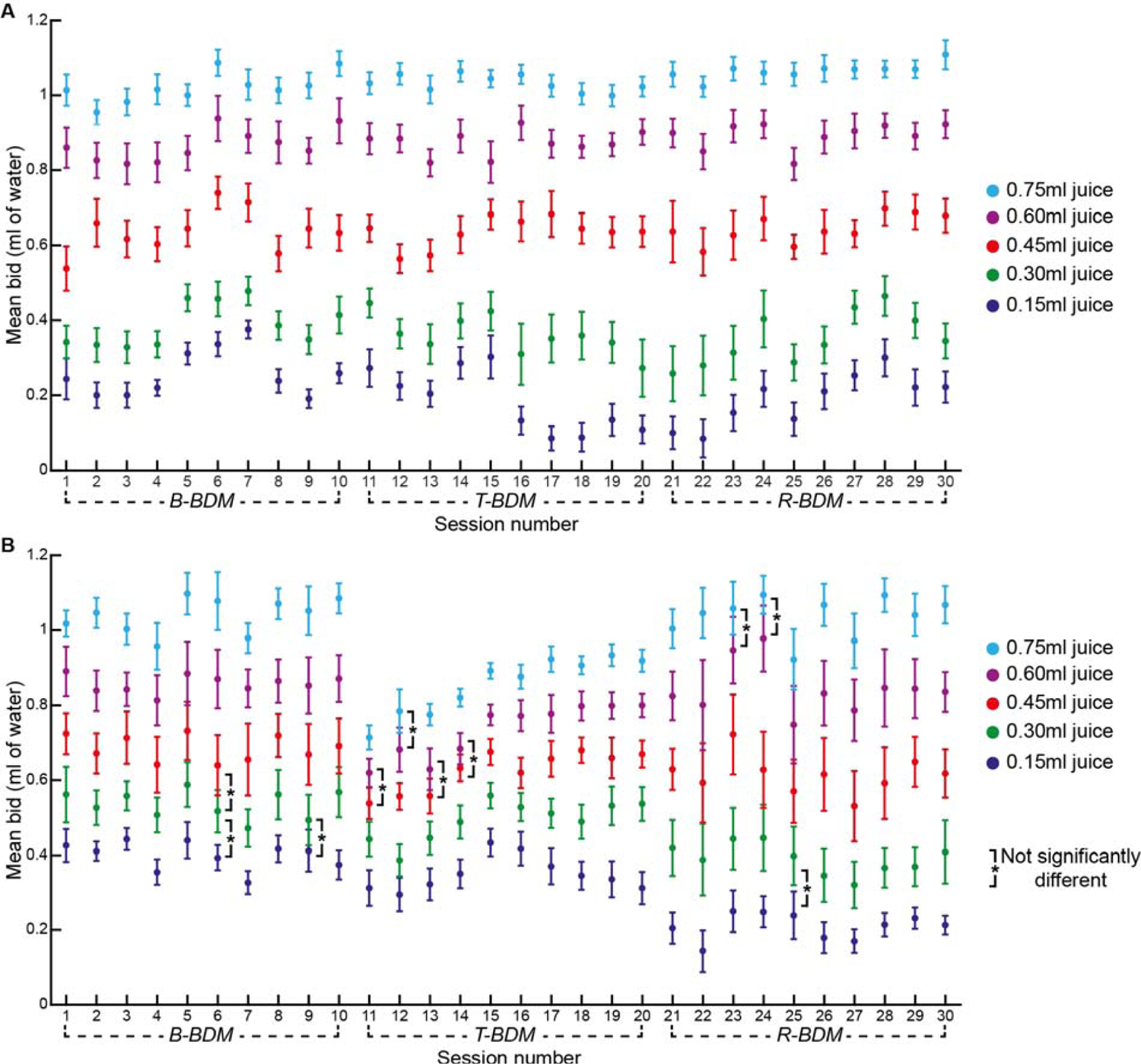
BDM bids in individual sessions. For tabulated numbers, see Extended Data Table 5-1. For development of differential bidding in early BDM task versions, see Extended Data Figs. 5-1 and 5-2. Sessions are numbered chronologically. ***A*,** Monkey A. All mean bids for each of the five juice volumes differed significantly in all 30 sessions. Error bars are 95% confidence intervals of the mean. In sessions 1-10 the bid cursor started at the bottom of the budget bar (B-BDM); for sessions 11-20 the cursor started at the top of the budget bar (T-BDM); and for sessions 21-30 the cursor started at a random position on the budget bar (R-BDM). Each session was composed of 200 correct trials. ***B*,** Monkey B. Mean bids differed significantly in 21 of the 30 sessions. In 8 sessions (1 B-BDMs; 4 T-BDMs; 3-RBDMs) the mean bids for two juice volumes differed insignificantly. In session 6 (B-BDM), the mean bid for the 0.30 ml juice differed insignificantly from those of either the 0.15 ml or 0.45 ml juice volumes. * in brackets indicates insignificant difference of mean bids after Bonferroni correction for multiple comparisons (α = 0.05).

Both animals consistently placed monotonically increasing bids for the five juice juice volumes (Fig. 6*A, B*). This positive monotonic relationship between bids and juice volume was significant in each of the 30 BDM sessions for both monkeys (Monkey A, Spearman Rho = 0.91 ± 0.02; mean ± SD; Monkey B, Spearman Rho = 0.81 ± 0.05; all *P* < 0.05; Table 1). Thus, the animals appropriately ranked the five juice volumes, and thus stated increasing subjective value for increasing juice volumes. More refined analyses in terms of juice utility revealed by the bids would have required estimation of the specific utility functions for the water the animals ‘paid’ for obtaining the desired juice (see Methods). Nevertheless, to substantiate the results from the Spearman analysis on the relationship between bids and juice volume, we found better fits with a linear function compared to nonlinear power and Prelec functions, thus confirming the bidding monotonicity (Extended Data Table 6-2).

**Figure 6.**
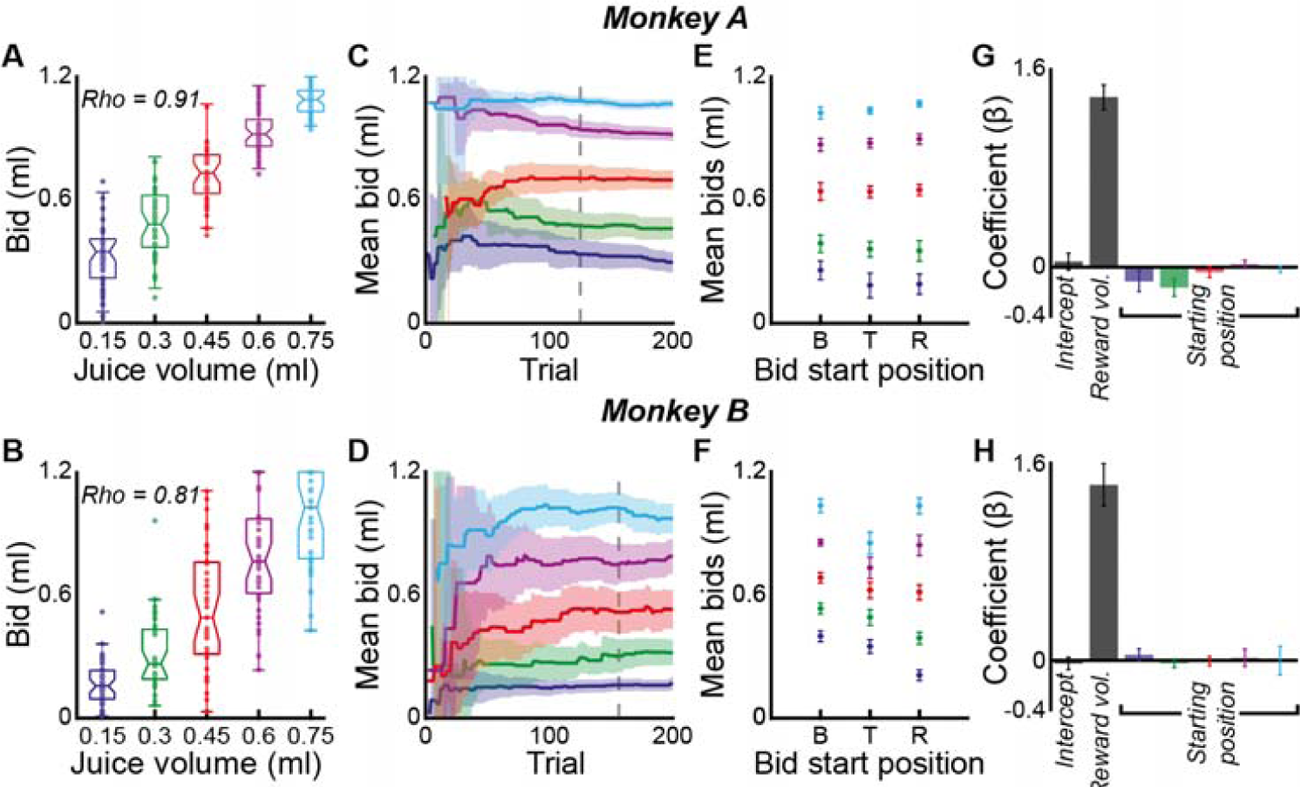
Increasing BDM bids with increasing juice volume, irrespective of bid cursor starting position. Top panels show session 28 for Monkey A, and bottom panels show session 27 is shown for monkey B (sessions chosen as Rho for these was closest to mean Rho across all sessions for each monkey). For tabulated numbers, see Extended Data Table 6-1. ***A, B*,** Monotonic increase of bids with juice volume in single sessions. Boxplots center lines show the median and notches show 95% confidence intervals of the median, boxplot edges mark interquartile range. Colors for juice volumes apply to all panels. ***C, D*,** Development of differential bidding across consecutive trials (same sessions as shown in A and B). Mean bids for all juice volumes became significantly different by trial 114 (Monkey A) and 170 (Monkey B) (*P* < 0.05, Bonferroni corrected t-test; grey dashed lines). Solid lines show mean bids, shaded areas show 95% confidence intervals. ***E, F*,** Similar discrimination of juice volumes by bids irrespective of bottom (B), top (T) or random (R) starting position (means of mean bids across all 10 sessions (*N* = 2,000 trials in each monkey) for each starting position). ***G, H*,** Mean beta coefficients from regression on juice volume and random starting position of bid cursor, for all five juice volumes (all 10 sessions in each monkey; *N* = 2,000 trials in each monkey) (Eq. 2). Bids varied significantly with cursor starting position only for the two smallest juice volumes with Monkey A (***G***: maroon, green). Error bars: 95% confidence intervals of the mean.

Moreover, whenever the animals achieved complete separation of all bids, they also achieved this before the end of the 200 correct trials that constituted a single testing session. On average, Monkey A needed 105.7 ± 38.4 trials (n = 30 sessions), and Monkey B needed 148 ± 30.1 trials (n = 21 sessions) to achieve complete separation of bids (Fig. 6*C, D*).

We applied stringent and conservative criteria to detect adequate separation of bids. We labelled the trial by which this was achieved as the 10^th^ trial in a row for which all mean bids had been statistically significantly different (alpha level of 0.05; with Bonferroni correction for multiple comparisons; see Materials and Methods: Statistical Analysis of BDM bids); at this point bids had been consistently well separated for some time.

Furthermore, we tested the animals with 5 distinct rewards in each session. In many tasks animals discriminate only between 2 or 3 different rewards. To approximate how well the BDM might be used to discriminate smaller sets of rewards, we performed the above analysis but taking combinations of 2 rewards, 3 rewards, and 4 rewards only, ignoring data for other rewards. For each session we again found the first trial by which mean bids for these rewards could be statistically discriminated in each session (Extended Data Fig. 6-1), revealing an expected linear relationship between the number of rewards tested and the point at which all means differed significantly for 10 consecutive trials.

Thus, the BDM could be used with fewer rewards to reliably distinguish bids using fewer trials. To further characterize the relationship between bid differentiation and number of bids for a given reward, we next looked at the first *N* trials for each of the 5 rewards for every session and used only these to perform Spearman’s rank correlation of bids on reward volumes, as well as one-way ANOVA and post-hoc t-tests, to identify significant differences in bids across the 5 juice volumes. Extended Data Fig. 6-2 shows the mean Spearman’s Rho (and *P* - value) across sessions and the mean number of significantly different pairwise comparisons between all 5 reward volumes (and their mean *P* - values) for every value of N between 2 and 20.

These analyses show that a significant positive monotonic correlation between bids and reward volumes was achieved in as few as 2 trials for each of the 5 rewards for both monkeys (Extended Data Fig. 6-2*A, B*). However far more trials were required to increase the number of significantly different pairwise comparisons of mean bids between the 5 different reward volumes (Extended Data Fig. 6-2*C, D*), reflecting the greater difficulty of separating noisy bids for 5 different rewards. Indeed, as suggested by the preceding analysis of smaller reward sets, reducing the number of rewards in the task would counter this.

Beyond such a significant positive correlation between bids and reward volumes, we were interested in how quickly monkeys could correctly rank all rewards with their bids. Within each session, both monkeys’ bids ranked all 5 rewards according to their reward volumes long before the end of the session. For Monkey A, this was typically achieved by trial 18.5 ± 11.8, with a significantly positive correlation between bids and reward volumes at this point (Spearman’s Rho = 0.87 ± 0.08). Similarly, Monkey B typically required only 19.6 ± 12.4 trials to achieve this (Spearman’s Rho = 0.74 ± 0.14).

Thus, the monkeys were both consistent in their ranking of rewards and in the precision of their bidding such that bids reliably reflected preferences and distinct subjective values for different rewards relatively early in each session, and within a single session of testing. These results demonstrate that monkeys were able to use the BDM to truthfully express reward values.

### Development of final BDM task

We used several successive steps to train both animals in the BDM task. First, they learned to associate different fractals on a computer monitor with different juice volumes. Then they learned to associate the budget bar on the computer monitor with different volumes of water. We also accustomed them to the sequential delivery of the water budget and the offered juice. They then learned to use a joystick in order to move the bid cursor and receive the different outcomes (win/loss) depending on the position of the computer bids relative to their own.

We then introduced the monkeys to various preliminary BDM task versions, using essentially similar types of fractal stimuli for juices but different volumes of juice reward and different volumes of water budget. We limited initially the reward volume in each trial such that the animal completed as many trials as possible on a test day. In earlier, reduced versions of the task with only three juice volumes and a lower budget volume, the monkeys ordered their bids according to their preferences, but their bids were inconsistent and poorly differentiated (Extended Data Fig. 5-1). We reasoned that while the relative cost of deviating from the optimal bid is unchanged by changing the budget volume, the absolute cost of a given deviation in terms of distance on the monitor, or movement of the joystick, is increased when larger rewards are on offer (Extended Data Fig. 5-2). With successively larger volumes of juice and water, bidding behavior improved, both in terms of correlation strength between bids and juice magnitude, as measured by Spearman rank correlation, and in terms of separation of bids for different juice volumes. For example, in an earlier task version with 0.6 ml of water as budget, Monkey A’s mean Spearman Rho for the correlation between bids and juice magnitude was 0.46 ± 0.085, compared to 0.91 ± 0.02 in the final task.

Similarly, for Monkey B, using 0.9 ml of water as the budget resulted in a mean Spearman Rho of 0.31 ± 0.26 for this correlation, compared to 0.81 ± 0.05 in the final BDM version. The larger volume limited the daily total trial numbers to 200.

Due to time constraints in testing earlier versions of the task, we changed several parameters at once (including juice type, magnitude and timing of stimulus presentation and reward delivery) and were unable to implement each change alone followed by a significant period of testing. This made it difficult to attribute any improvement in performance to a single parameter change or manipulation of the task structure. Nevertheless, the improvements we observed using larger budget volumes in these unstructured preliminary tests guided our approach in using a larger budget volume for the final BDM task presented here.

#### Control for action effects

Both monkeys’ bids discriminated all juice volumes regardless of initial cursor position (Fig. 6*E, F*). Two-way unbalanced ANOVAs with factors of juice volume, bid cursor starting condition and their interaction demonstrated a highly significant effect of juice volume on the monkeys’ bids (Monkey A: F_4,5985_ = 6889.46, *P* < 0.001, ω^2^= 0.82; Monkey B: F_4,5985_ = 2353.17, *P* < 0.001, ω^2^= 0.58) (Extended Data Table 6-1). Bid cursor starting position had a smaller but still significant effect (Monkey A: F_2,5985_ = 7.18, *P* < 0.001, ω^2^ = 3.67 × 10^-4^; Monkey B: F_2,5985_ = 148.94, *P* < 0.001, ω^2^ = 0.018). The interaction between juice volume and starting position was also significant (Monkey key A: F_8,5985_ = 13.55, *P* < 0.001, ω^2^ = 3 × 10^-3^; Monkey B: F_8,5985_ = 55.86, *P* < 0.001, ω^2^ = 0.027). Thus, while the starting position of t^ω^he bidding cursor affected bidding to some extent, differential bidding for juice volume remained significant irrespective of the starting position and the juice volume accounted for most of the variance in bids.

To interrogate the influence of motor contingencies on bidding more closely, we further analysed the bids from the 10 sessions in which the cursor’s starting position varied randomly (R-BDM). In these sessions, the bid cursor appeared at a random vertical position, requiring different joystick movements to achieve the same bids from trial to trial. We regressed the monkeys’ bids on both juice volume (JV) and cursor starting position (SP) for each of the 10 R-BDM sessions individually, such that:

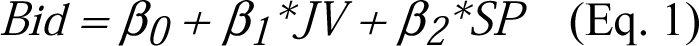

Across these 10 sessions, we found that the monkeys’ bids varied significantly with the juice volume (Monkey A: β_1_ = 1.53 ± 0.12; Monkey B: β_1_ = 1.40 ± 0.10), with a far smaller effect of the cursor’s starting position for monkey A (β_2_ was significantly smaller than zero; β_2_ = −0.06 ± 0.05) but with no effect of starting position for monkey B (β_2_ = 0.01 ± 0.05). To investigate for any variable effect of starting position with different juice volumes, we then performed a regression of the monkeys’ bids on both juice volume (JV) and cursor starting position separately for each of the five juice volumes (SP_JV=Xml_), such that:

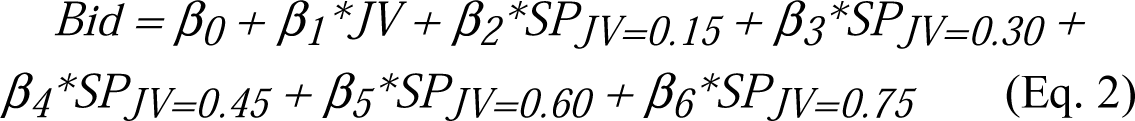

The results from this analysis confirmed a small but significant effect of starting position for the two smallest juice volumes for Monkey A (β_2_ = −0.11 ± 0.12, mean ± SD; β_3_ = −0.17 ± 0.10), but none of the position coefficients differed significantly from zero for Monkey B (Fig. 6*G, H*). For Monkey A this may have reflected reduced motivation to bid precisely on trials that promised lower juice volumes. Nevertheless, juice volume had a far greater influence on the final bid than cursor starting position, for both monkeys (Monkey A: β_1_ = 1.38 ± 0.14; Monkey B: β_1_ = 1.42 ± 0.24).

These results suggest that the animals were not merely responding with greater vigor to larger juice volumes, or just learning conditioned motor responses. Their bids reflected the value of the juice irrespective of the specifics of the required joystick movement.

### Comparison of BDM bids and BC values

#### Values inferred using the BC task

While the positive monotonic relationship of BDM bids to juice volumes in both monkeys suggests systematic value estimation, it is important to know whether these results were specific for the BDM mechanism or were independent of the eliciting mechanism. A different eliciting mechanism would also provide independent estimates for assessing optimality in BDM bidding. Therefore, we compared the values inferred from BDM bids with estimates from a conventional value eliciting method commonly used in animals. (Note that while the study’s goal was to assess subjective juice value in single BDM trials, comparison with value estimation by conventional binary choice required repeated measures.)

We implemented a binary choice (BC) task with repeated trials that used the same options, visual stimuli and juice and water outcomes as the BDM task and differed only in the choice aspect (Fig. 7*A*). Option 1 contained a bundle comprised of one of the five juice volumes and a varying, partial water amount, equivalent to the outcome when winning the BDM. Option 2 contained the full water budget, equivalent to the outcome when losing the BDM. Thus, when choosing the juice-water bundle, the monkey forewent some of the full water budget to obtain the juice (like winning in the BDM); when choosing the other option, they received the full water budget but no juice (like losing in the BDM). We performed 10 of these BC sessions, and each session consisted of 200 trials. In each session every reward volume appeared in one of 10 possible bundles (i.e. with 10 different possible volumes of water in the bundle), and each of these combinations was repeated 4 times per session, such that there were 40 trials per reward volume in each session, for a total of 200 trials.

**Figure 7.**
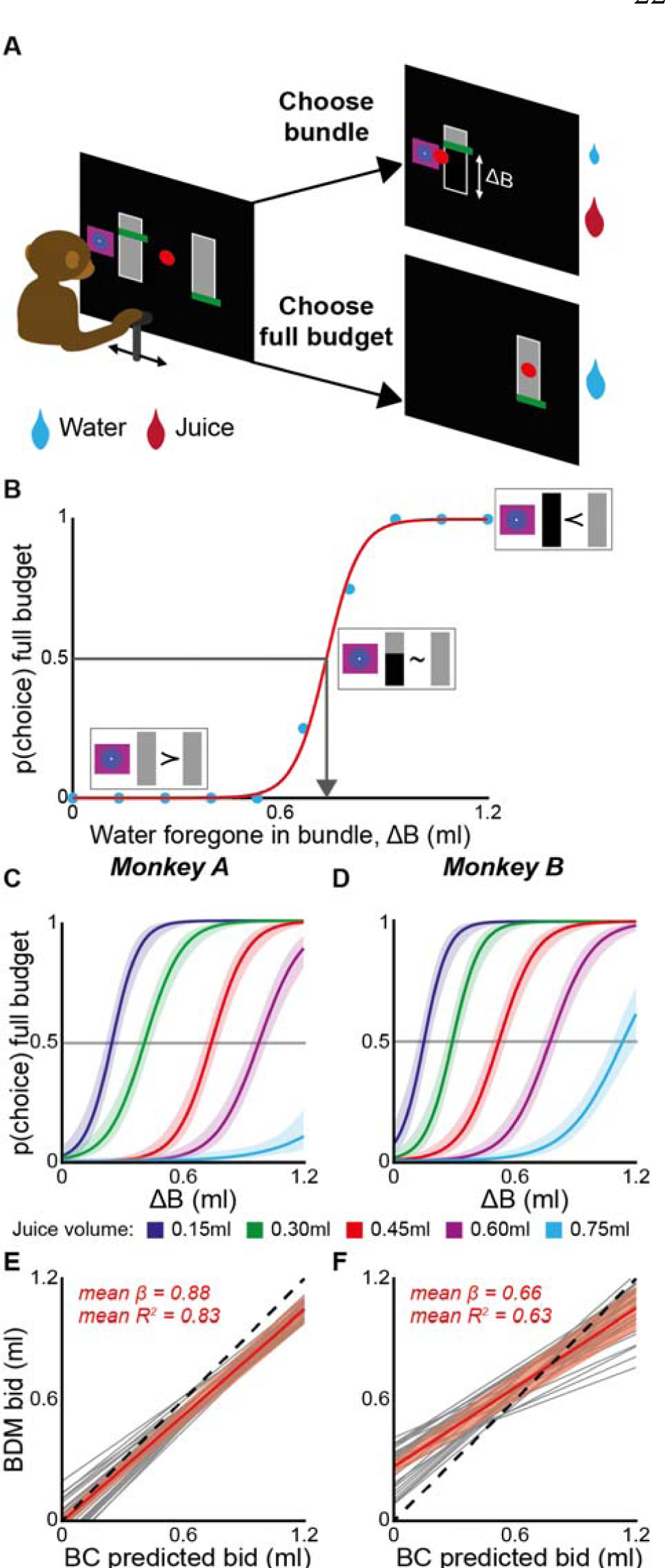
Mechanism independence: comparison with value estimation in Binary Choice (BC) task. *A*, BC task. Choice between [bundle of specific juice volume (fractal) combined with a specific water volume (grey area above green line) (option 1)] and [full water budget (full grey vertical rectangle) (option 2)]. The monkey indicated its choice by moving a horizontal joystick-driven red dot onto the preferred option. At left, the grey rectangle below the green line (bundle, option 1) represented the water foregone (ΔB) from the full budget and is blackened after the monkey’s choice (see ‘Choose bundle’ at right). Left and right option positions alternated pseudorandomly. ***B*,** Psychophysical value estimation of juice value in the currency of water during BC. Decrease of water in option 1 increased the choice probability of option 2. At choice indifference (P (choice) = 0.5, grey line), the water foregone in the bundle (ΔB) indicated the value of the juice volume in units of ml of water. A logistic regression (red) was fitted to the monkey’s choices (blue). More preferred (≻); indifferent (∼); less preferred (≺). ***C, D*,** BC value estimates for each of the five juice volumes used in the BDM. Choices were pooled across all 10 BC sessions (n = 2000 trials) for each monkey. Shaded areas are 95% confidence intervals of the fitted logistic function. See Extended Data Fig. 7-1 for individual sessions and mean bids, and Extended Data Table 7-1 for BDM and BC values. ***E, F*,** Regression of monkeys’ bids on the best bid as predicted by the BC task. The best bid was equal to the BC task value estimate, or, the maximum bid of 1.2 ml, whichever was smaller. The identity line is dashed. The mean from the fits of all individual sessions is shown in red, and the red shaded area shows the 95% confidence interval; fits for individual sessions are shown in grey.

**Figure 8.**
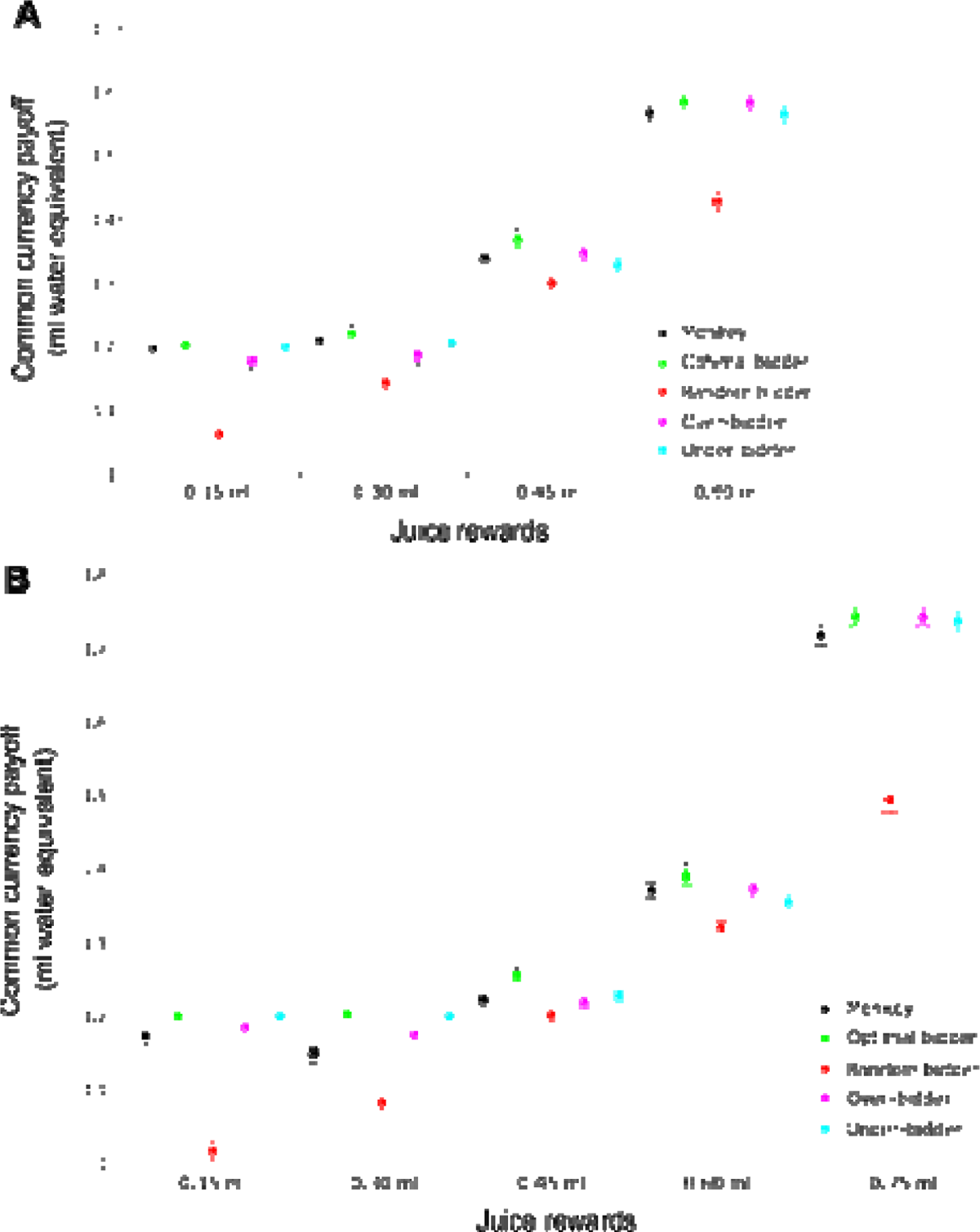
Optimality of BDM bids. Average individual trial payoffs are shown for each monkey (***A****, **B***) and expressed in equivalent water value, for the actual monkey bidder (black) and the simulated optimal bidder (green), random bidder (red), over-bidder (magenta) and under-bidder (cyan). Significant differences of pairwise comparisons are shown with an asterisk and lines indicating which pairs had a significant difference. Payoffs for the random bidder were always significantly lower than for all other bidders. Error bars show 95% confidence intervals.

**Figure 9.**
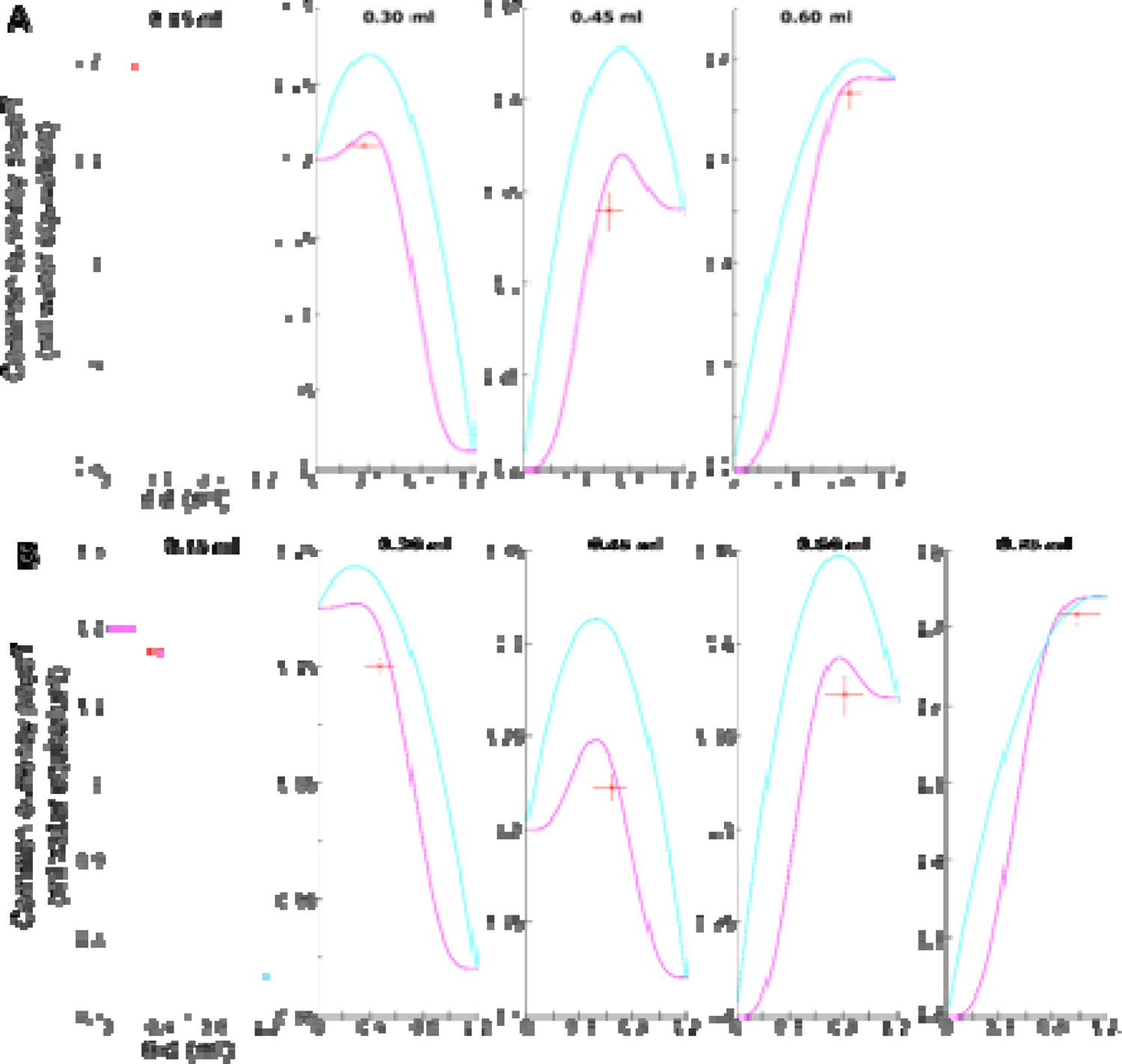
Modeled expected payoffs of all bids. The magenta curves show the expected payoffs (y-axis; ml of water) for the currently used Beta(4,4) distribution (magenta) for generating the BDM computer bids; the cyan curves show the analogous payoffs for a uniform distribution (cyan). Each panel shows the payoffs for a separate reward (labelled at the top of each panel). The monkey’s mean bid and mean payoff intersect at the red cross. The horizontal red line shows the interquartile range of bids, and the vertical line shows the 95% confidence intervals for the mean payoff. Data are shown for both monkeys (***A, B***).

Choice preference among the two options varied systematically (Fig. 7*B*). The monkeys showed little choice of the full water budget (option 2) when the alternative juice-water bundle (option 1) contained substantial water amounts in addition to the juice; apparently the slight loss in water volume was overcompensated in value by the added juice (Fig. 7*B* left). Choice of the full water budget increased gradually with more water foregone in the juice-water bundle (ΔB against the full water budget). At some specific volume of water foregone, the monkey preferred the full water budget as much as the juice-water bundle (Fig. 7*B* center; P (choice) = 0.5; choice indifference). At this point, the juice together with the remaining water was valued as much as the full water budget alone; hence the juice compensated fully for the water foregone and was valued as much as that water volume (ΔB). Thus, the value of the juice can be expressed on a common currency basis in ml of water volume foregone at choice indifference (ΔB). In this way, psychophysics allowed us to estimate the value for each specific juice volume being tested.

### Mechanism independence

In both monkeys, the choice indifference points in the BC task followed the same rank order as the BDM bids for the five juice volumes (Fig. 7*C, D*; see Extended Data Fig. 7-1*A-C* for individual sessions and Extended Data Table 7-1 for BDM and BC values).

We performed 5 BC sessions before and 5 after the 30 BDM sessions and found the BC estimates of value were stable across this period of BDM testing (Extended Data Fig. 7-1*E, F*). We therefore pooled choices across all 10 sessions of the BC task to infer an estimate of value for each juice reward in terms of water volume across sessions. Thus, each value estimate we used in subsequent analyses was inferred from 400 pooled trials of the BC task (10 sessions, with each reward presented 40 times per session). Accordingly, Pearson correlation coefficients between the bids elicited across all 30 BDM sessions and the value estimates from all 10 BC sessions were high (Monkey A: 0.91 ± 0.02; Monkey B: 0.79 ± 0.05). To confirm these results and provide more detail, we performed a least-squares regression of BDM bids on the values estimated by the BC task, such that:

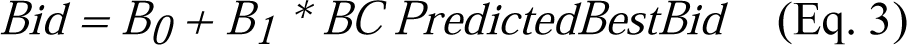

The PredictedBestBid inferred from performance in the BC task is equal to the water value of the chosen option in the BC task, except when the BC value is greater than the maximum possible bid of 1.2 ml of water, in which case the best possible bid is equal to 1.2 ml, as was the case for the 0.75 ml reward for Monkey A. An optimal bidder’s BDM bids should perfectly reflect the value for the commodity (B_1_ = 1) without any bias in bidding (B_0_ = 0) (the value may, for example, be modulated by the mental and/or motor effort of placing a bid). BDM bids correlated closely with the BC estimates for both Monkey A (mean B_1_ = 0.88 ± 0.09, and mean R^2^ = 0.83 ± 0.03) and Monkey B (mean B_1_ = 0.66 ± 0.15, mean R^2^ = 0.63 ± 0.08) (Fig. 7*E, F*). Monkey A did not have any significant bidding bias (B_0_ = 0 ± 0.09), but monkey B had a significant bias which accounted for overbidding for low juice volumes (B_0_ = 0.27 ± 0.10).

In showing good correlations between single BDM bids and conventional binary stochastic choices with both numerical methods, these data suggest that value estimation by BDM largely reflects the same underlying value also elicited by the BC task and is not simply determined by the specific elicitation method. Previous experiments using the BDM method have not attempted to compare values thus elicited with those elicited by any other means and have simply taken the BDM validity as a given. Here, we show a high degree of concordance between the BDM task and the standard binary choice method of value elicitation in animals. Thus, the BDM in monkeys appears to provide a novel and valid mechanism for estimating subjective economic value in animal subjects.

### Optimality of bidding in the BDM

The incentive compatibility of the BDM rests on the notion that bidders benefit most by stating their value accurately for a given reward (Material and Methods: Optimal BDM Strategy).

However, unlike human subjects in the BDM, animals cannot be made explicitly aware of the optimal strategy for maximising reward value. Instead, they adjust their bidding behavior according to the experienced outcome. Further, performance in the BDM provides less intuitive assessments due to its second-price nature, and BDM outcomes are risky because they depend on the computer bid drawn from a fully specified probability distribution. By contrast, stimuli in the BC task display the options in a direct and explicit manner, and the animal gets exactly what it has chosen.

Therefore, we used the economic values estimated in the BC task to assess optimal bidding for each juice volume. Specifically, the optimal bid is equal to the PredictedBestBid stated above and is derived from the combined value of both the juice and the water budget, as expressed in common currency units of ml of water.

To assess the optimality of BDM bidding, we compared each monkey’s payoffs to those of hypothetical bidders: an optimal bidder, who always bids the BC value for each juice volume according to the best BDM strategy; a random bidder, whose bids are drawn from the same uniform distribution for all juice volumes; and over-/under-bidders, consistently bidding ±0.2 ml for each reward (Material and Methods: Simulated Bidding). These simulated bidders faced the same 6,000 juice presentations and computer bids as the monkeys did across 30 sessions of BDM testing (200 correct trials each). We used pairwise t-tests, corrected for multiple comparisons (α = 0.5), to detect significant differences in payoffs between the different types of bidders for each juice reward.

The differences between the average per-trial payoffs for the simulated optimal bidder and for monkey A were small across all rewards (Fig. 8*A*). These ranged from a difference of only 0.0059 ± 0.03 ml for the 0.15 ml reward, to a difference of 0.028 ± 0.063 ml for the 0.45 ml reward, however, the optimal bidder only secured a significantly greater payoff than Monkey A for the 0.3 ml and 0.45 ml rewards (all p < 0.01). The systematic over- and under-bidders performed similarly to Monkey A, and never secured a significantly greater payoff, but the over-bidder had a significantly smaller average per-trial payoff for the 0.15 ml and 0.3 ml rewards (all p < 0.01). The random bidder consistently secured significantly smaller payoffs (all p < 0.01) than all other bidders (all results summarised in Extended Data Table 8-1; payoffs could not be computed for the 0.75 ml juice for this animal as the value for this volume was above the possible bidding range).

For Monkey B (Fig. 8*B*), the absolute differences in average per-trial payoff between the optimal bidder and Monkey B were again small and ranged from 0.018 ± 0.052 ml for the 0.6 ml reward to a difference of 0.053 ± 0.11 ml for the 0.3 ml reward; the optimal bidder secured a significantly greater payoff for the 0.15 ml, 0.3 ml, and 0.45 ml rewards (all p < 0.01). Monkey B secured significantly smaller payoffs than the under-bidder for the 0.15 ml and 0.3 ml rewards, and less than the over-bidder for the 0.3 ml reward (all p < 0.01). Again, the random bidder secured significantly smaller payoffs than all other bidders for all rewards (all p < 0.01; Extended Data Table 8-1).

These small differences in average per-trial payoff were comparable to the juice delivery system’s error due to the variability of droplet size (SD = 0.06 ml per trial; Material and Methods: Juice-delivery error), which may have been too small to be perceived by the monkeys. Moreover, bidders that deviate as much as 0.2 ml (1/6^th^ of the range of possible bids, from 0 ml to 1.2 ml) from their value are evidently still rewarded with payoffs of similar magnitude to an optimal bidder. Together, these factors limit performance in the BDM task, as difficulty discriminating small differences in payoffs limits learning of the optimal bid, and the cognitive and motor effort costs of placing precise bids may outweigh the benefits of doing so.

To better appreciate the relationship between bids and payoffs, we can calculate the payoff for every possible bid, and for each reward, given the water values found in the BC task. The expected profit,, for a given bid, ***b***, and value, ***v***, against a computer drawing a price, ***p***, from a probability density function, ***f(p)*** with a cumulative distribution function, ***F(p)***, is given by the following equation (Lusk and Shogren, 2007):

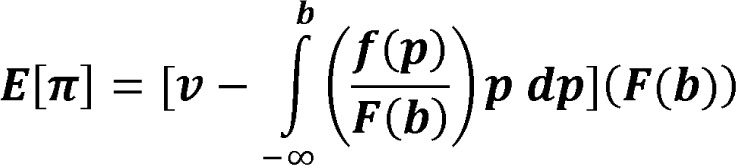

The overall expected payoff on a given trial is the starting budget in addition to the above ‘profit’ (which may be negative). The maximum expected payoff is always found where the bid is equal to the value; elsewhere, the relationship between a given bid and expected payoff is determined by the distribution from which computer bids are drawn. Fig. 9 shows the modeled expected payoffs given the Beta(4,4) distribution we used (magenta) for all possible bids and for each reward separately for both monkeys. It is apparent that payoffs may be similar over a wide range of bids close to the reward’s value; for example, for the 0.15 ml reward for monkey A, all bids lower than 0.5 ml grant an expected payoff between 1.15 ml and 1.2 ml. Thus, the 0.2 ml over- and under-bidding of simulated bidders are unlikely to experience dramatically different payoffs when compared to an optimal bidder. The red crosses in Fig. 9 demonstrate that the monkey’s actual bidding fell within this optimal window.

Fig. 9 also shows the payoffs given by bids drawn from a modeled uniform distribution (cyan). We used a Beta(4,4) distribution, which has a lower expected payoff for all bids and thus may help to increase the number of trials in neuronal recording experiments. Moreover, a Beta(4,4) distribution is associated with a steeper decline in payoff with less accurate bidding (Extended Data Fig. 9-1) than the uniform distribution across most of the bidding range; the decline is particularly prominent in the centre of the bidding range where the monkeys suboptimally tended to cluster their bids in an earlier task version (Extended Data Fig. 5-1*A*). In conclusion, the Beta(4,4) distribution results in a steeper relationship between bids and payoffs in this range as compared to a uniform distribution; this characteristic of our Beta distribution increases the cost of deviating from the optimal bid and therefore encourages learning of the optimal strategy and hence optimal bidding.

## Discussion

This study shows that monkeys can truthfully report their internal, subjective economic value of rewards in individual trials by placing bids in a BDM auction-like bidding mechanism. The monkeys reliably and systematically ranked their preferences over five juice volumes, and their bids correlated well with values inferred from choices in an equivalent BC task, in keeping with the theoretical ‘incentive compatibility’ of both methods. While the monkeys’ bids were noisy, they nevertheless achieved a level of performance that approximated that of a simulated optimal bidder and well exceeded that of a random bidder. One of the difficulties in comparing behavior between BDM and BC tasks may lie in the inadvertent, unintended but unavoidable inclusion of ambiguity and risk in BDM that is not a necessary component in BC tasks but likely affects performance and adds uncertainty attitude to the choices. These experiments contribute a novel method of value elicitation for research on economic decision-making in monkeys and show the capacity of monkeys to perform auction-like bidding in resemblance to human behavior; indeed, human bids are also noisy, and unlike the results presented here, have not previously been compared to values elicited in BC tasks.

Subjective value elicitation in animals has thus far relied upon the use of repeated choices (Platt and Glimcher 1999; Padoa-Schioppa and Assad 2006; Kobayashi and Schultz 2008), inferring an average, single subjective value from dozens of decisions that are performed with some degree of stochasticity (whether in making the decision or executing it). However, decisions are typically made in single instances, weighing subjective values on a moment-to-moment basis, and usually have immediately tangible consequences. Accordingly, human experimental economics research considers decisions and assesses values in individual trials.

To better understand the underlying processes in animals and achieve a greater degree of concordance in research across species, we need methods that elicit values within single choices. Moreover, while the BDM requires the decision-maker to generate a bid, and thus acceptable cost, in a continuous manner, BC tasks involve a more constrained discrete choice between only two possibilities. Thus, the BDM directly elicits a value, while the BC task only allows to derive value secondarily from measurable choices. It is unclear whether the valuation processes during choice between two options, and the underlying neuronal mechanisms, are the same as, or different to, those required to actively state a value for oneself as in BDM.

These features of the BDM make it worth investigating in animals, but they also present unique limitations and challenges. Our BDM task required more extensive training than an equivalent BC task. First, because the placement of bids on a continuous scale is a complex motor task compared to simple left/right choices. And second, because the BDM’s second-price nature means that deviating from the optimal strategy is not strongly punished; for example, many bids that are too high still result in the bidder paying less from the budget than they think the item is worth, as long as the computer bid is lower than their value for that item. Thus, on any given BDM trial, over- or under-bidding may not lead to a loss relative to bidding one’s value (although it would lead to a loss in the long run). Conversely, choosing the less desired option in the BC task always leads to a definite loss relative to the preferred option. Therefore, up to a point, decision costs may even outweigh the benefits of precise bidding in the BDM; introducing an element of decision noise which may not be present, or as impactful, in BC tasks.

The BDM provides a rich decision-making scenario which brings the scope of animal studies of decision-making closer to those in humans. At the same time, the complexity of the BDM offers new opportunities for study. Monkeys, at a basic level of reward function, have a globally similar brain organisation to humans, and the feasibility of a behavioral task used frequently in humans could provide unprecedented information about the role of single reward and decision neurons in auction-like mechanisms.

Using a theoretically equivalent method to verify the values elicited by the BDM can only provide a best approximation, as any two differing methods are likely to be confounded by psychological differences. On the other hand, these differences may be the subject of future research. For example, while ambiguity has been introduced into BC tasks by occluding information in the two choice options (Hayden et al., 2010), it is unclear to what extent animals are averse to ambiguity when they have some degree of control over this. In the BDM the monkey could reduce ambiguity by its own actions; by increasing or reducing its bid, the monkey makes the outcome of a trial more certain. Moreover, the effects of ambiguity on valuation in the BDM could be interrogated by changing the variance of the computer bid distribution, as reducing the variance about the mean increases the certainty of a given computer bid and reduces the ambiguity in outcomes for monkey bids that are distant from this mean computer bid.

It is not enough to interrogate the activity of neurons in the presence of rewards; rather, for understanding reward processing, animals should reveal their preferences by making choices (Platt and Glimcher, 1999; Stauffer et al., 2014). Besides conventional BC tasks, experimenters may now benefit from eliciting truthful valuation by different means while examining neuronal processes underlying those choices. It would also be interesting to see the extent to which the existing data from conventional BC tasks depend on their specific eliciting mechanism. For example, neurons encoding action-specific reward values have been identified in the striatum (Samejima et al., 2005), but it is not known whether these reward values were specific to the decision rules and contexts in which they were elicited.

Because monkeys in the BDM report values on a trial-by-trial basis, the task provides a closer temporal relationship to the activity of single neurons and allows to capture trial-by-trial fluctuations in value not visible with choices across many trials. The suitability of BDM bidding for neuronal recordings in monkeys is further supported by the current finding that action only affects reward valuation to a very limited extent; different actions, as required by different bidding start positions, did not substantially affect reward valuation in our task. Thus, the BDM task as presently designed might be an alternative to standard BC tasks when searching for fine-grained single-trial neuronal variations related to rapid spontaneous changes of subjective reward value.

The reported meaningful BDM performance was obtained with substantial experimental constraints. Natural wildlife does not prepare monkeys for explicitly stating their values against some odds, even though animals always need to make some form of commitment to satisfy their needs. Their good performance highlights their adaptive cognitive abilities. One contributing factor might have been the seating for several hours in a primate chair that may have helped the animals to focus onto the task. Our use of tangible, ecologically relevant and familiar liquids may have also been beneficial, whereas it is unclear how the monkeys would have performed when bidding for more abstract items, such as tokens used in neurophysiological BC experiments (Seo and Lee 2009). Thus, future work may help to delineate the conditions in which rhesus monkeys are able to successfully perform a BDM task.

The primate BDM makes the link to human studies in several ways. Apparently, the relative closeness in cognitive functions between human and monkey would not only explain their successful BDM bidding but also allow for more direct comparisons with human neuroimaging studies, as the BDM is commonly used in experimental work (Plassmann et al., 2007; Chib et al., 2009; Harris et al., 2011; Tang et al., 2014; Tyson-Carr et al., 2018) as well as in consumer economics (Linder et al., 2010). Whereas human neuroimaging provides a larger overview of brain processes, single-neuron electrophysiology provides better cellular resolution for distinction of valuation functions in different neuron types. In this way, the current BDM data provide both an evolutionary and methodological link between the two primate species.

## Author contributions

AAM and WS designed the study, AAM performed experiments, analyzed data and constructed figures, AAM and WS wrote the paper.

## ACKNOWLEDGEMENTS

We thank Aled David and Christina Thompson for animal and technical support and Alexander Pastor-Bernier, Raymundo Baez-Mendoza, Philipe Bujold, Fabian Grabenhorst, Armin Lak, William R. Stauffer, Louis Nerurkar, Juliet Griffin, John O. Ledyard and Charles R. Plott for discussions on the design of this experiment. This study was supported by Wellcome Trust (WT 095495, WT 204811), European Research Council (ERC; 293549) and US National Institutes of Mental Health (NIMH) Caltech Conte Center (P50MH094258). This research was funded in part by the Wellcome Trust [206207/Z/17/Z]. For the purpose of Open Access, the author has applied a CC BY public copyright licence to any Author Accepted Manuscript version arising from this submission.

## Extended Data Figures and Tables

**Extended Data Figure 5-1.**
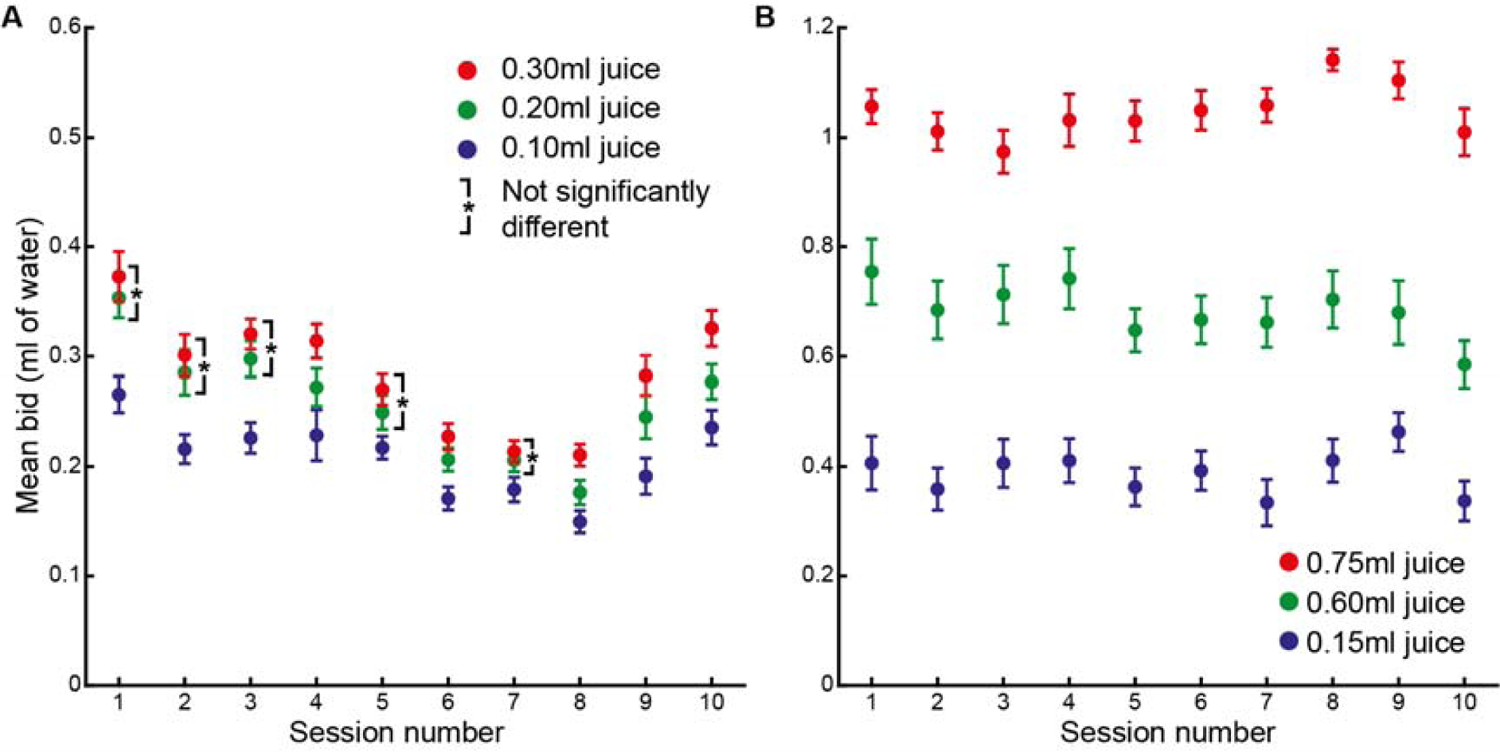
Performance in early BDM task versions. Juice volumes were selected from performance in a preceding binary choice task such that their values covered a wide range of possible bids. All bids started at the bottom. Error bars show 95% confidence intervals of the mean. Data shown for Monkey A. ***A*,**Early version of BDM task with small water budget volume (0.6 ml) and 3 small juice volumes to be bid for. Small volumes maximised the number of trials in each session before satiety set in; however, bids were not well differentiated, and the correlation between juice volumes and bids was weaker than in later task versions (mean Spearman Rho = 0.45±0.25). Asterisks indicate insignificantly varying mean bids after Bonferroni correction for multiple comparisons (α=0.05). ***B*,** We hypothesised that an increase in the water budget and juice volumes would lead to more careful bidding as the absolute losses for a given deviation in terms of distance from the optimal bid would be increased. We therefore doubled the water budget volume to 1.2 ml and used larger juice volumes, such that the range of juice reward values covered this wider range of possible bids. This led to a marked performance improvement, with mean bids for all juice volumes being significantly different to one another in every session. Moreover, the correlation between juice volumes and bids was markedly and consistently stronger than in the lower budget volume version of the task shown in A (mean Spearman Rho = 0.80±0.03).

**Extended Data Figure 5-2.**
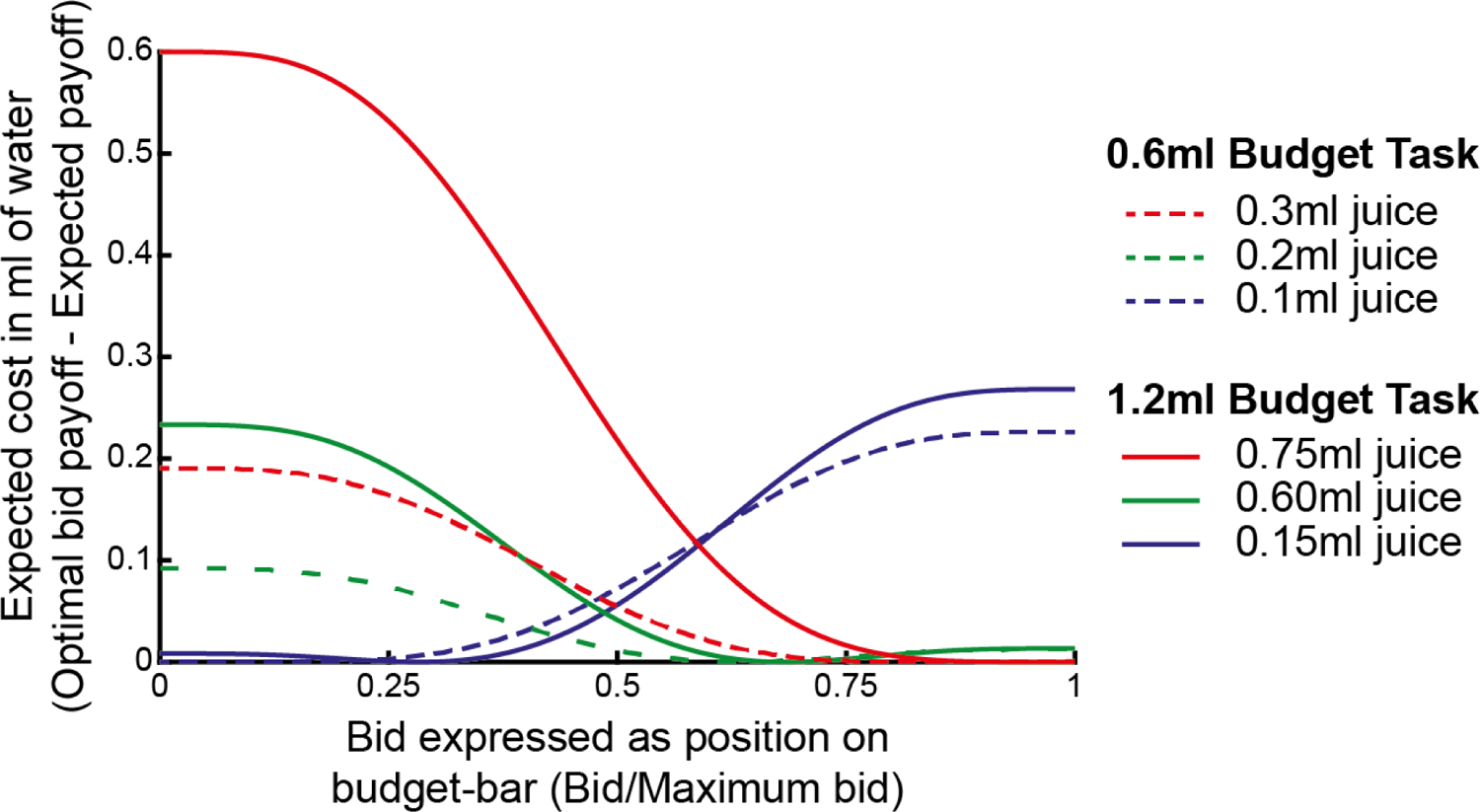
Increasing expected suboptimal bidding cost with increasing juice and water budget. The optimal BDM bid is equal to the value of the juice volume being bid for and will lead to the highest expected payoff compared to all other bids. The lower expected payoff of other bids constitutes an expected cost relative to the optimal bid. In the two BDM payoff settings shown in Fig. S4, the 0.3 ml and 0.75 ml, 0.2 ml and 0.6 ml, and 0.1 ml and 0.15 ml juice volumes elicited optimal bids that were similarly positioned on the 0.6 ml and 1.2 ml budget bars used in each task, respectively. This can be seen by the fact that the minimum costs for these pairs of juice volumes are at similar positions on the budget bar. For a given deviation of the final bid in terms of distance on the budget bar, the cost is higher in the 1.2 ml budget task than in the 0.6 ml budget task. This effect is more pronounced the further bids are away from the centre of the bidding range, because the mean computer bid was at the centre of this range. Moreover, the effect is exaggerated for lower bids for higher juice volumes, as the cost of losing a higher juice volume by bidding less than its value is greater.

**Extended Data Figure 6-1.**
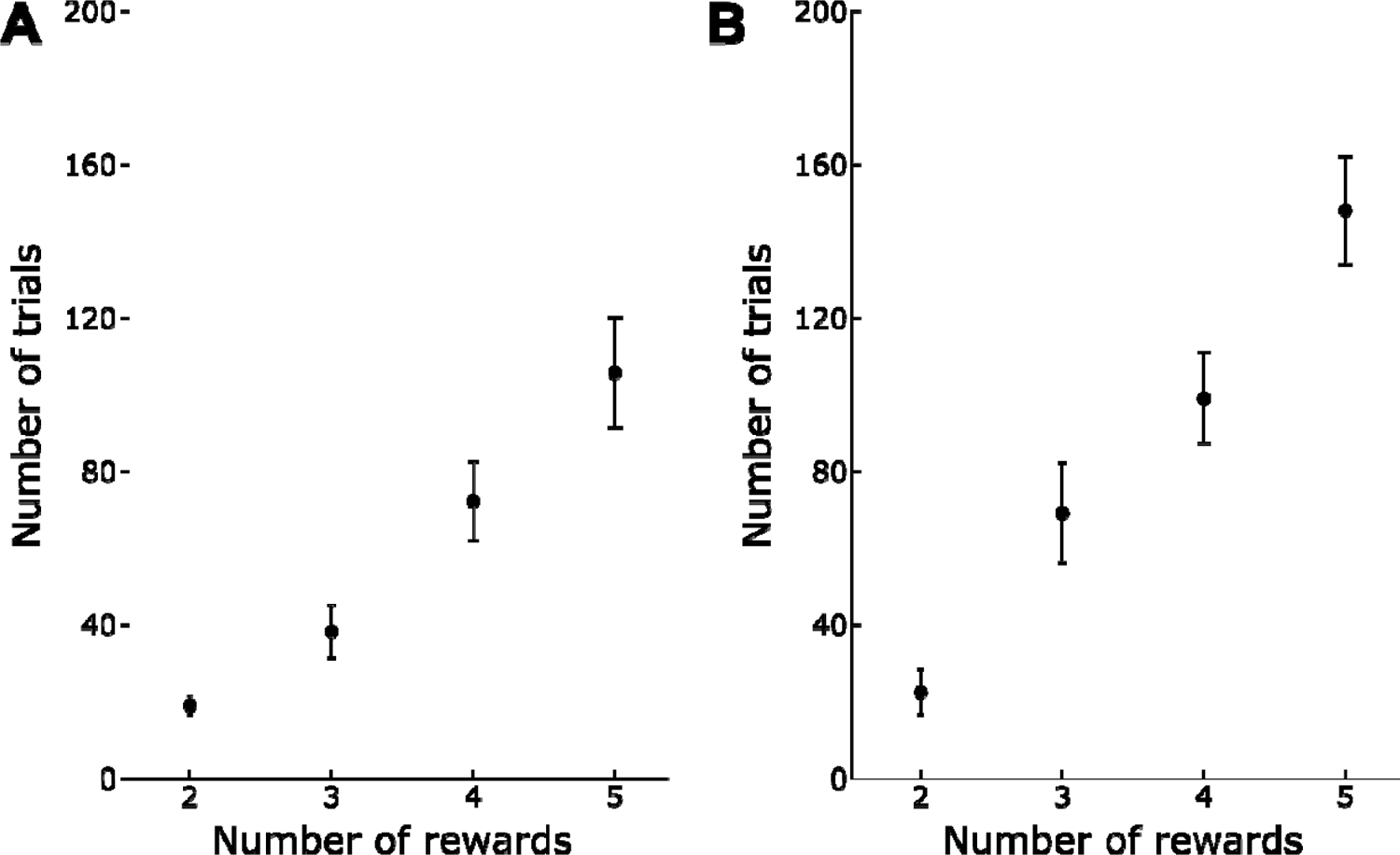
Increasing the number of rewards increases the number of trials required to achieve consistent statistical separation of bids. We generated different combinations of 2, 3, or 4 rewards and found the first trial by which bids for these different rewards differed statistically significantly to one another for 10 consecutive trials. Here we show the mean trial number, across sessions, by which this was achieved for 2, 3, and 4 rewards for both monkeys *A* and *B*). The data using all 5 rewards are shown for comparison. There was a significant positive relationship between the number of rewards being bid for and the number of trials required to achieve such separation of bids (Pearson’s R = 0.79, p = 6.17 × 10-27 for monkey A; Pearson’s R = 0.88, p = 7.58 × 10-27 for monkey B). Error-bars are 95% confidence intervals of the mean.

**Extended Data Figure 6-2.**
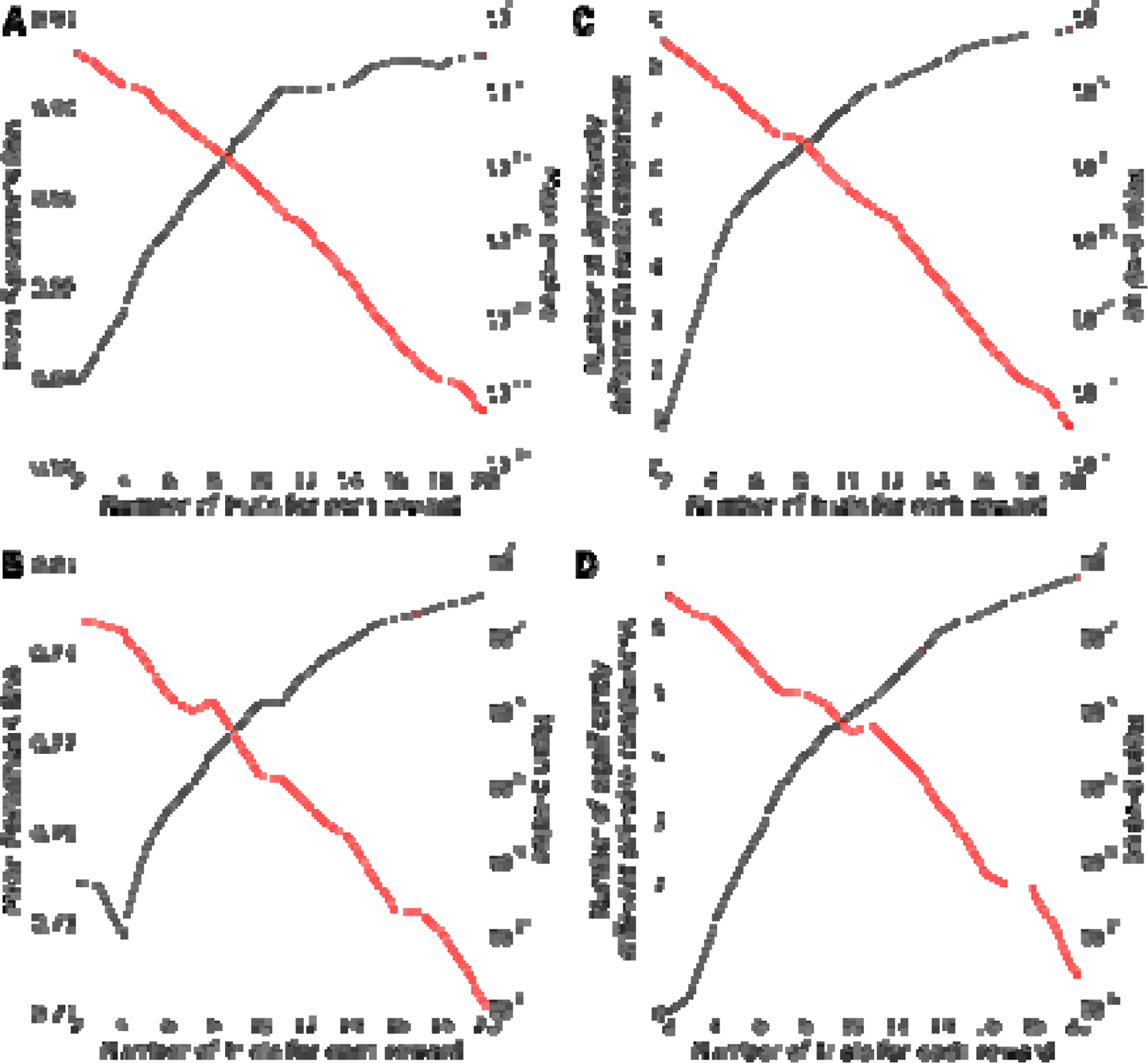
Relationship between number of trials for each reward and BDM performance. *A, B*, Spearman’s Rho for correlations between bids and reward volumes, taking only the first *N* trials for each reward (x-axis). Monkeys A (***A***) and B (***C***) required only 2 bids for each reward to demonstrate statistically significant positive monotonic relationship between reward volume and bids (Spearman’s rank correlation). ***C, D,*** Results from t-tests between bids for all rewards on the first *N* trials for each reward (x-axis), and mean number of significantly different pairwise comparisons at each value of *N* across all sessions. Increasing the number of trials for each reward increased the number of significantly different pairwise comparisons, suggesting more reliable differentiation of rewards using BDM bids. The requirement of complete separation of means for all rewards was more stringent than the requirement of a significant positive relationship (Spearman’s rank) or some difference in means (one-way ANOVA); even with 20 trials per reward, the monkeys, on average, did not achieve significance in all possible pairwise comparisons (for 5 rewards there are 10 possible pairwise comparisons). Mean p-values for Spearman’s rank correlation (***A, B***) and t-tests (***C, D***) are shown in red and relative to the right-sided y-axis of each panel, using a logarithmic scale. Dashed line shows the p-value for comparisons. Α = 0.05, corrected for multiple

**Extended Data Figure 7-1.**
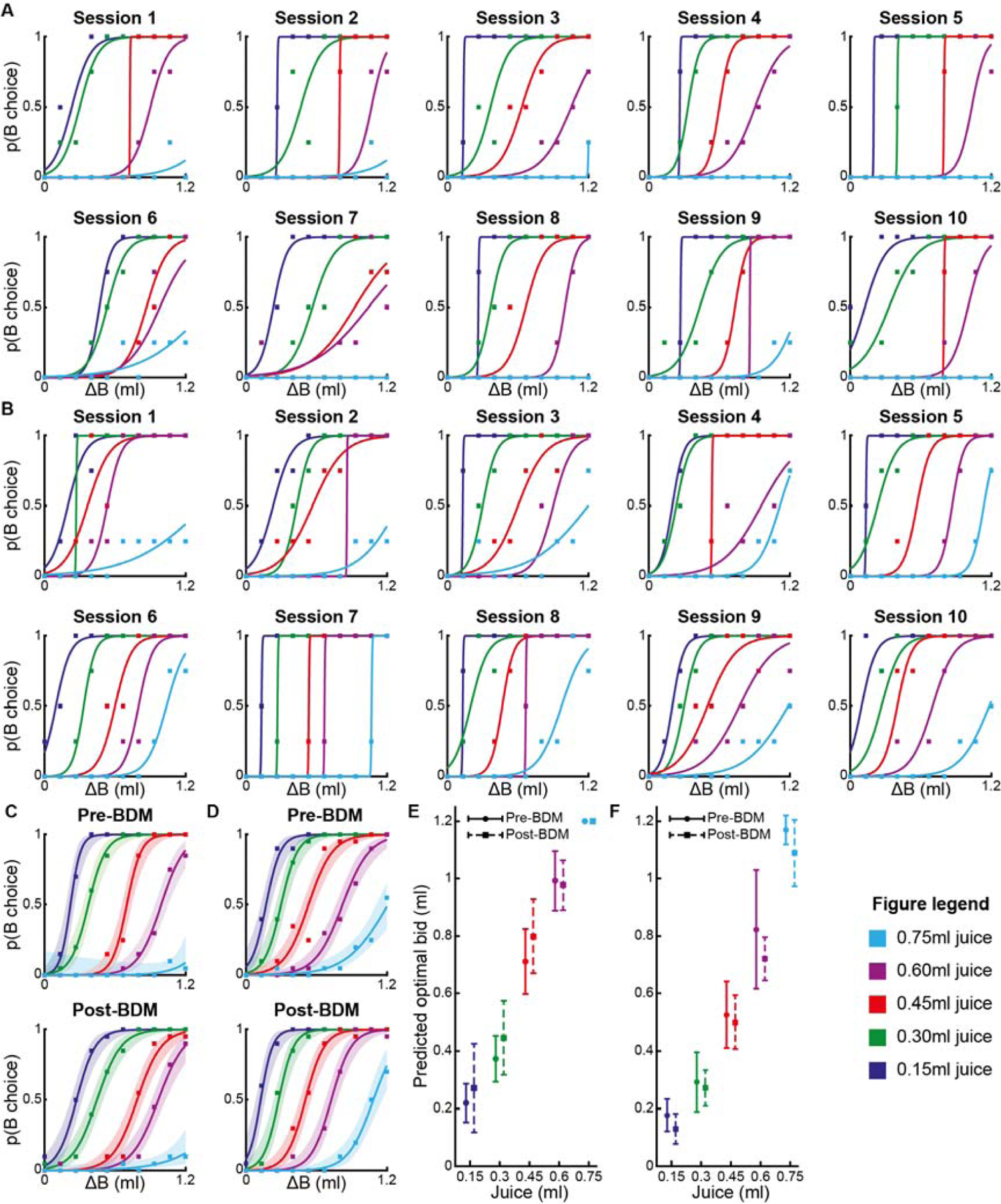
Choice probabilities in Binary Choice task, and pre- and post-BDM comparison. *A*, Lines of best fit for logistic regression of choice probability of full budget, p(B choice), on water volume foregone in each bundle (ΔB). Monkey A, individual sessions. ***B*,** as A, but Monkey B, individual sessions. ***C, D*,** As ***A*** and ***B***, respectively, but pooled from 5 session before BDM (Pre-BDM) and 5 sessions after all 30 BDM sessions (Post-BDM). ***E, F*,** Comparison of mean predicted optimal bids for each juice volume from 5 Binary Choice task sessions before BDM (Pre-BDM; solid lines) and 5 sessions after BDM (Post-BDM; dotted lines), for Monkeys A and B, respectively. Changes in predicted optimal bid for any of the juice volumes was insignificant for either monkey (two-tailed Student t-tests, all *P* > 0.05). Error bars are 95% confidence intervals of the mean.

**Extended Data Figure 9-1.**
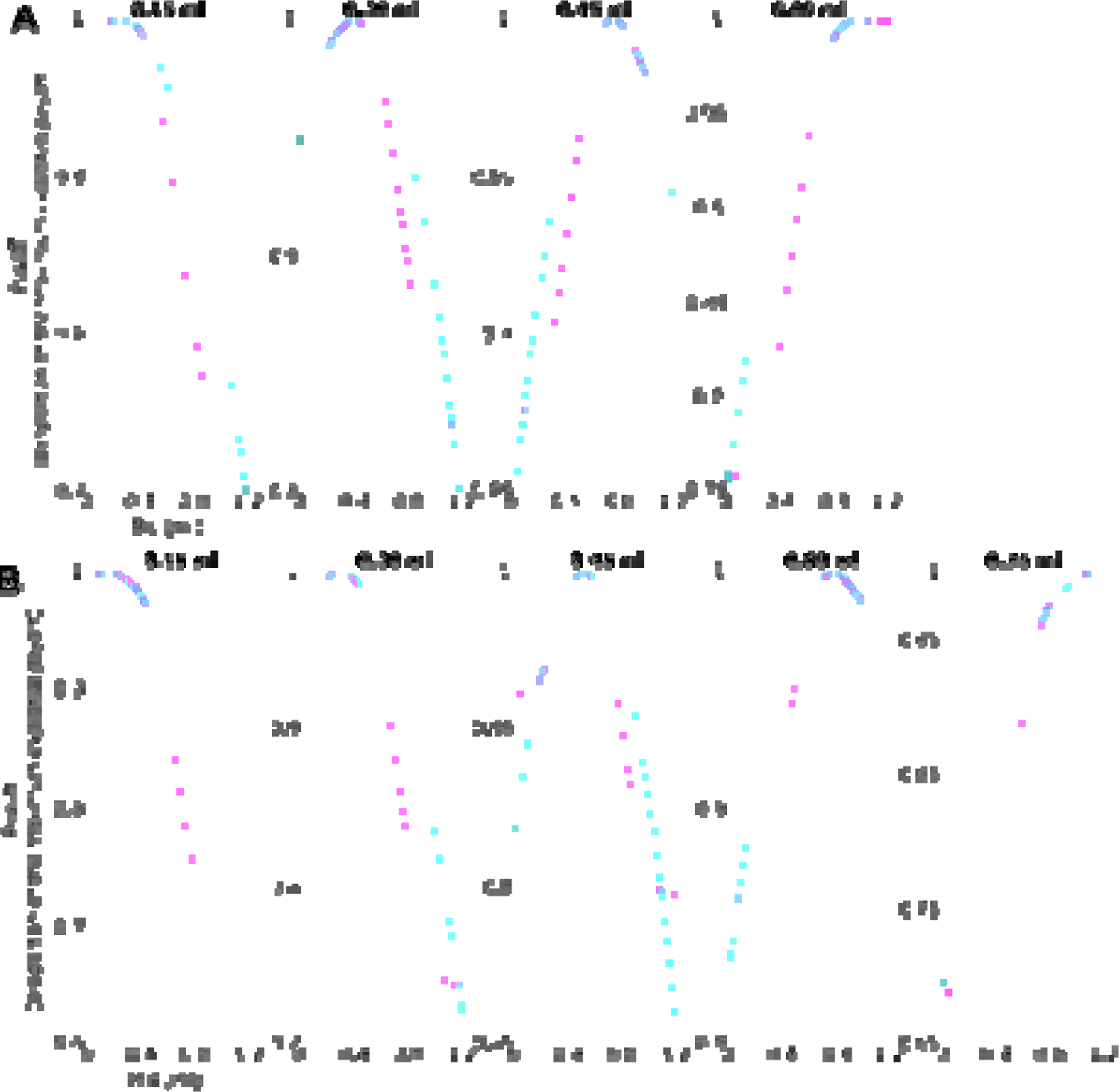
**Differences in payoff decline between different modeled computer bid distributions**. In the Beta(4,4) distribution (magenta), the payoff declines more rapidly with increasing distance from optimal bid compared to the uniform distributions (cyan). This steeper decline encourages accurate bidding. The payoff for every bid is expressed as a proportion of the maximum expected payoff for that reward. The Beta(4,4) distribution can be seen to be relatively less rewarding than the uniform distribution for a given deviation away from the optimal bid (where the maximum payoff is found and is equal to 1 here) in the centre of the bidding range, whereas the uniform distribution payoffs are relatively lower at the extremes of the bidding range. In an earlier version of the task (Extended Data Fig. 5-1*A*) the monkey was observed to cluster bids towards the centre of the bidding range. A Beta(4,4) distribution was used to encourage optimal bidding by increasing the monkey’s relative costs of deviating from their value in the central part of the bidding range. Payoffs are shown for both monkeys (***A****, **B***). The payoffs shown here are expressed as proportion of the maximum payoffs; the absolute payoffs are shown in Fig. 9.

**Extended Data Table 2-1.** Error data for Monkey A. Shown are the total number of errors and percentage of trials that resulted in an error (each session consisted of 200 correct trials). Also shown are the number and percentage of trials in which there was no movement of the joystick, as well as the incidence of each type of error and the percentage of errors in each session attributed to each error type.

| Type | Session | Errors | No Joystick movement | No Hold | No Choice | Not centred |
| --- | --- | --- | --- | --- | --- | --- |
| **B-BDM** | **1** | 21 (9.5%) | 20 (95.2%) | 18 (85.7%) | 2 (9.5%) | 1 (4.8%) |
|  | **2** | 36 (15.3%) | 34 (94.4%) | 23 (63.9%) | 12 (33.3%) | 1 (2.8%) |
|  | **3** | 29 (12.7%) | 29 (100%) | 22 (75.9%) | 5 (17.2%) | 2 (6.9%) |
|  | **4** | 27 (11.9%) | 27 (100%) | 21 (77.8%) | 5 (18.5%) | 1 (3.7%) |
|  | **5** | 52 (20.6%) | 52 (100%) | 42 (80.8%) | 10 (19.2%) | 0 (0%) |
|  | **6** | 31 (13.4%) | 31 (100%) | 18 (58.1%) | 9 (29.0%) | 4 (12.9%) |
|  | **7** | 97 (32.7%) | 96 (99.0%) | 15 (15.5%) | 5 (5.2%) | 77 (79.4%) |
|  | **8** | 15 (7.0%) | 15 (100%) | 14 (93.3%) | 1 (6.7%) | 0 (0%) |
|  | **9** | 10 (4.8%) | 9 (90.0%) | 0 (0%) | 10 (100%) | 0 (0%) |
|  | **10** | 2 (1.0%) | 0 (0%) | 0 (0%) | 2 (100%) | 0 (0%) |
| **T-BDM** | **11** | 8 (3.8%) | 7 (87.5%) | 2 (25.0%) | 5 (62.5%) | 1 (12.5%) |
|  | **12** | 6 (2.9%) | 5 (83.3%) | 0 (0%) | 6 (100%) | 0 (0%) |
|  | **13** | 22 (9.9%) | 22 (100%) | 1 (4.5%) | 21 (95.5%) | 0 (0%) |
|  | **14** | 5 (2.4%) | 5 (100%) | 0 (0%) | 5 (100%) | 0 (0%) |
|  | **15** | 14 (6.5%) | 14 (100%) | 0 (0%) | 14 (100%) | 0 (0%) |
|  | **16** | 5 (2.4%) | 4 (80.0%) | 0 (0%) | 5 (100%) | 0 (0%) |
|  | **17** | 7 (3.4%) | 7 (100%) | 0 (0%) | 6 (85.7%) | 1 (14.3%) |
|  | **18** | 8 (3.8%) | 7 (87.5%) | 0 (0%) | 8 (100%) | 0 (0%) |
|  | **19** | 1 (0.5%) | 1 (100%) | 0 (0%) | 0 (0%) | 1 (100%) |
|  | **20** | 3 (1.5%) | 3 (100%) | 0 (0%) | 2 (66.7%) | 1 (33.3%) |
| **R-BDM** | **21** | 2 (1.0%) | 1 (50.0%) | 1 (50.0%) | 1 (50.0%) | 0 (0%) |
|  | **22** | 5 (2.4%) | 1 (20.0%) | 1 (20.0%) | 4 (80.0%) | 0 (0%) |
|  | **23** | 3 (1.5%) | 1 (33.3%) | 0 (0%) | 2 (66.7%) | 1 (33.3%) |
|  | **24** | 12 (5.7%) | 0 (0%) | 0 (0%) | 12 (100%) | 0 (0%) |
|  | **25** | 1 (0.5%) | 1 (100%) | 0 (0%) | 0 (0%) | 1 (100.0%) |
|  | **26** | 1 (0.5%) | 1 (100%) | 0 (0%) | 0 (0%) | 1 (100%) |
|  | **27** | 3 (1.5%) | 2 (66.7%) | 1 (33.3%) | 1 (33.3%) | 1 (33.3%) |
|  | **28** | 6 (2.9%) | 4 (66.7%) | 1 (16.7%) | 2 (33.3%) | 3 (50.0%) |
|  | **29** | 1 (0.5%) | 1 (100%) | 0 (0%) | 0 (0%) | 1 (100%) |
|  | **30** | 0 (0%) | 0 (0%) | 0 (0%) | 0 (0%) | 0 (0%) |

**Extended Data Table 2-2.**
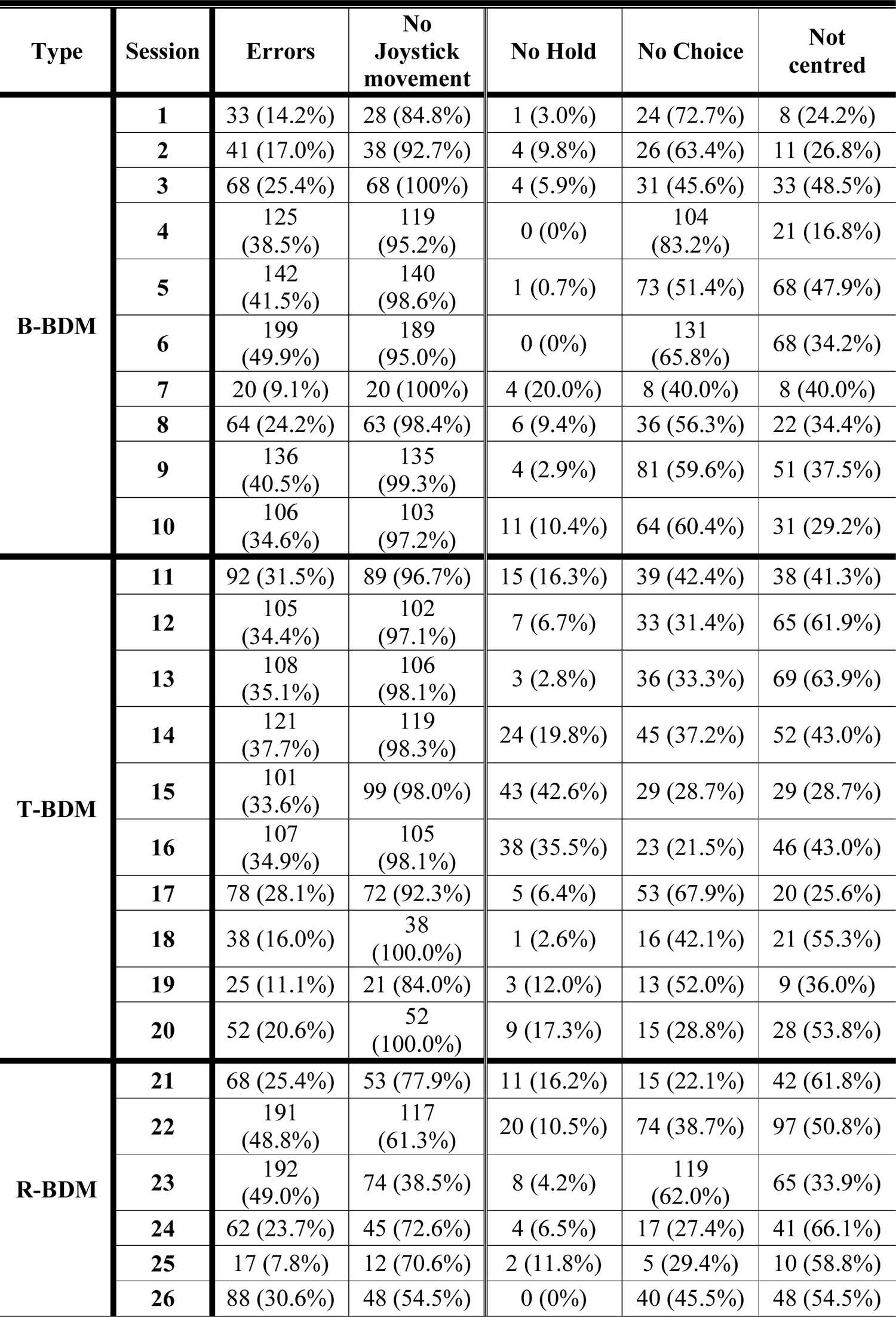
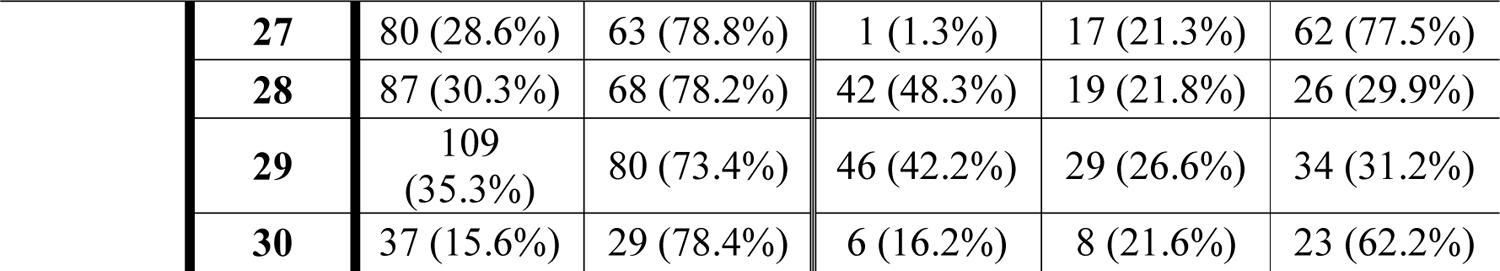
Error data for Monkey B. Shown are the total number of errors and percentage of trials that resulted in an error (each session consisted of 200 correct trials). Also shown are the number and percentage of trials in which there was no movement of the joystick, as well as the incidence of each type of error and the percentage of errors in each session attributed to each error type.

| Type | Session | Errors | No Joystick movement | No Hold | No Choice | Not centred |
| --- | --- | --- | --- | --- | --- | --- |
| **B-BDM** | **1** | 33 (14.2%) | 28 (84.8%) | 1 (3.0%) | 24 (72.7%) | 8 (24.2%) |
|  | **2** | 41 (17.0%) | 38 (92.7%) | 4 (9.8%) | 26 (63.4%) | 11 (26.8%) |
|  | **3** | 68 (25.4%) | 68 (100%) | 4 (5.9%) | 31 (45.6%) | 33 (48.5%) |
|  | **4** | 125 (38.5%) | 119 (95.2%) | 0 (0%) | 104 (83.2%) | 21 (16.8%) |
|  | **5** | 142 (41.5%) | 140 (98.6%) | 1 (0.7%) | 73 (51.4%) | 68 (47.9%) |
|  | **6** | 199 (49.9%) | 189 (95.0%) | 0 (0%) | 131 (65.8%) | 68 (34.2%) |
|  | **7** | 20 (9.1%) | 20 (100%) | 4 (20.0%) | 8 (40.0%) | 8 (40.0%) |
|  | **8** | 64 (24.2%) | 63 (98.4%) | 6 (9.4%) | 36 (56.3%) | 22 (34.4%) |
|  | **9** | 136 (40.5%) | 135 (99.3%) | 4 (2.9%) | 81 (59.6%) | 51 (37.5%) |
|  | **10** | 106 (34.6%) | 103 (97.2%) | 11 (10.4%) | 64 (60.4%) | 31 (29.2%) |
| **T-BDM** | **11** | 92 (31.5%) | 89 (96.7%) | 15 (16.3%) | 39 (42.4%) | 38 (41.3%) |
|  | **12** | 105 (34.4%) | 102 (97.1%) | 7 (6.7%) | 33 (31.4%) | 65 (61.9%) |
|  | **13** | 108 (35.1%) | 106 (98.1%) | 3 (2.8%) | 36 (33.3%) | 69 (63.9%) |
|  | **14** | 121 (37.7%) | 119 (98.3%) | 24 (19.8%) | 45 (37.2%) | 52 (43.0%) |
|  | **15** | 101 (33.6%) | 99 (98.0%) | 43 (42.6%) | 29 (28.7%) | 29 (28.7%) |
|  | **16** | 107 (34.9%) | 105 (98.1%) | 38 (35.5%) | 23 (21.5%) | 46 (43.0%) |
|  | **17** | 78 (28.1%) | 72 (92.3%) | 5 (6.4%) | 53 (67.9%) | 20 (25.6%) |
|  | **18** | 38 (16.0%) | 38 (100.0%) | 1 (2.6%) | 16 (42.1%) | 21 (55.3%) |
|  | **19** | 25 (11.1%) | 21 (84.0%) | 3 (12.0%) | 13 (52.0%) | 9 (36.0%) |
|  | **20** | 52 (20.6%) | 52 (100.0%) | 9 (17.3%) | 15 (28.8%) | 28 (53.8%) |
| **R-BDM** | **21** | 68 (25.4%) | 53 (77.9%) | 11 (16.2%) | 15 (22.1%) | 42 (61.8%) |
|  | **22** | 191 (48.8%) | 117 (61.3%) | 20 (10.5%) | 74 (38.7%) | 97 (50.8%) |
|  | **23** | 192 (49.0%) | 74 (38.5%) | 8 (4.2%) | 119 (62.0%) | 65 (33.9%) |
|  | **24** | 62 (23.7%) | 45 (72.6%) | 4 (6.5%) | 17 (27.4%) | 41 (66.1%) |
|  | **25** | 17 (7.8%) | 12 (70.6%) | 2 (11.8%) | 5 (29.4%) | 10 (58.8%) |
|  | **26** | 88 (30.6%) | 48 (54.5%) | 0 (0%) | 40 (45.5%) | 48 (54.5%) |

|  |  |  |  |  |  |
| --- | --- | --- | --- | --- | --- |
| **27** | 80 (28.6%) | 63 (78.8%) | 1 (1.3%) | 17 (21.3%) | 62 (77.5%) |
| **28** | 87 (30.3%) | 68 (78.2%) | 42 (48.3%) | 19 (21.8%) | 26 (29.9%) |
| **29** | 109<br>(35.3%) | 80 (73.4%) | 46 (42.2%) | 29 (26.6%) | 34 (31.2%) |
| **30** | 37 (15.6%) | 29 (78.4%) | 6 (16.2%) | 8 (21.6%) | 23 (62.2%) |

**Extended Data Table 5-1.**
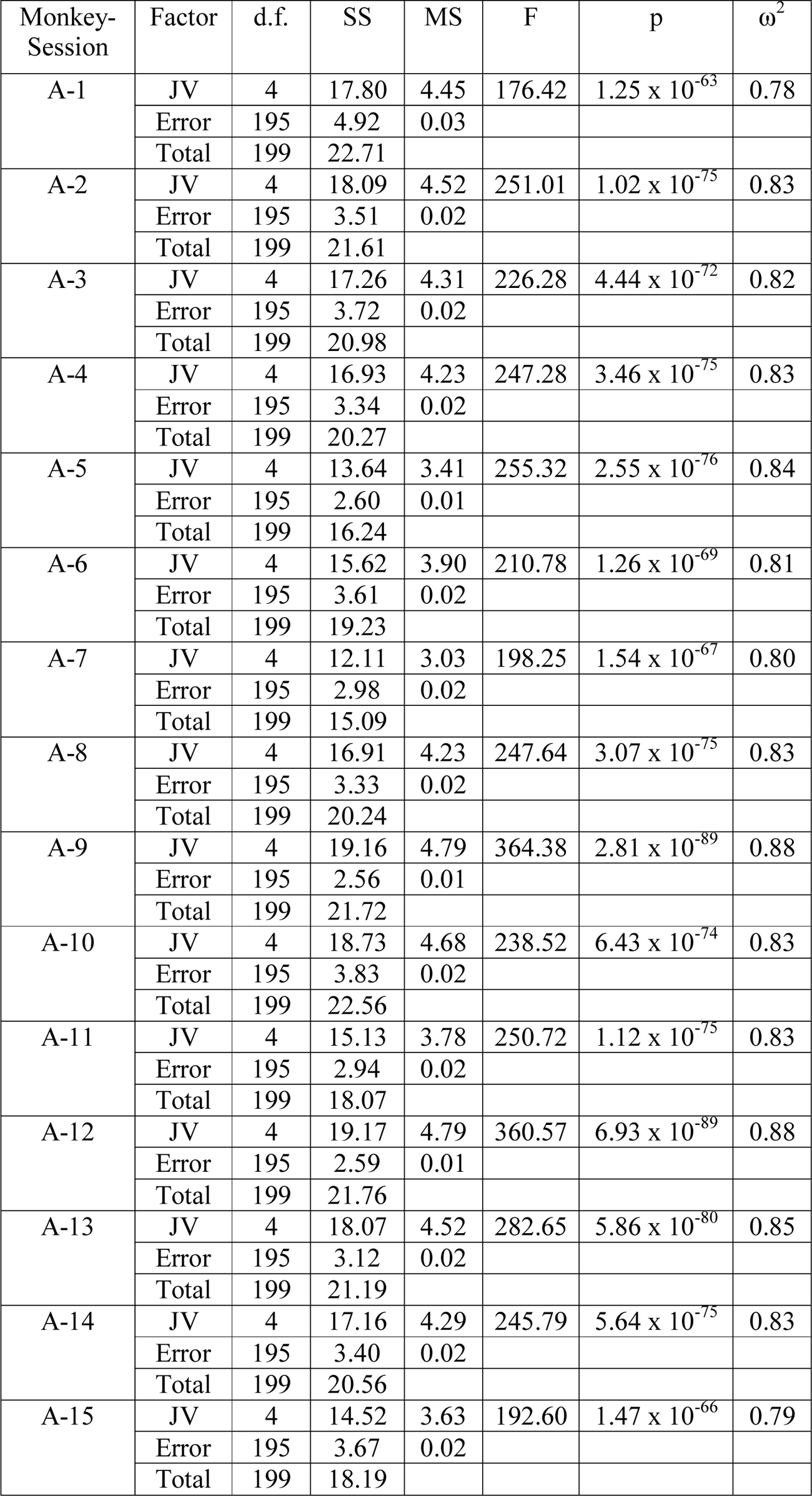
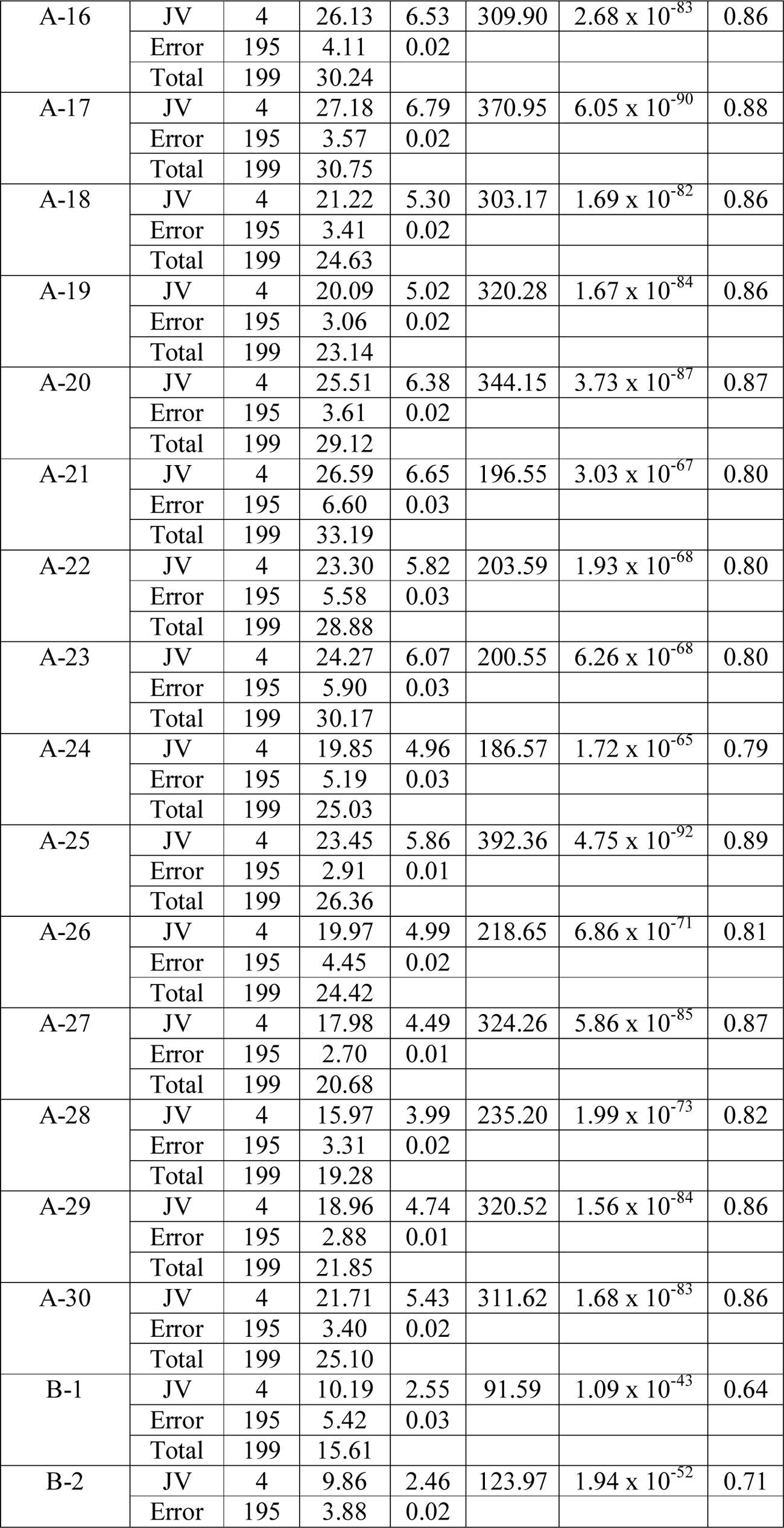
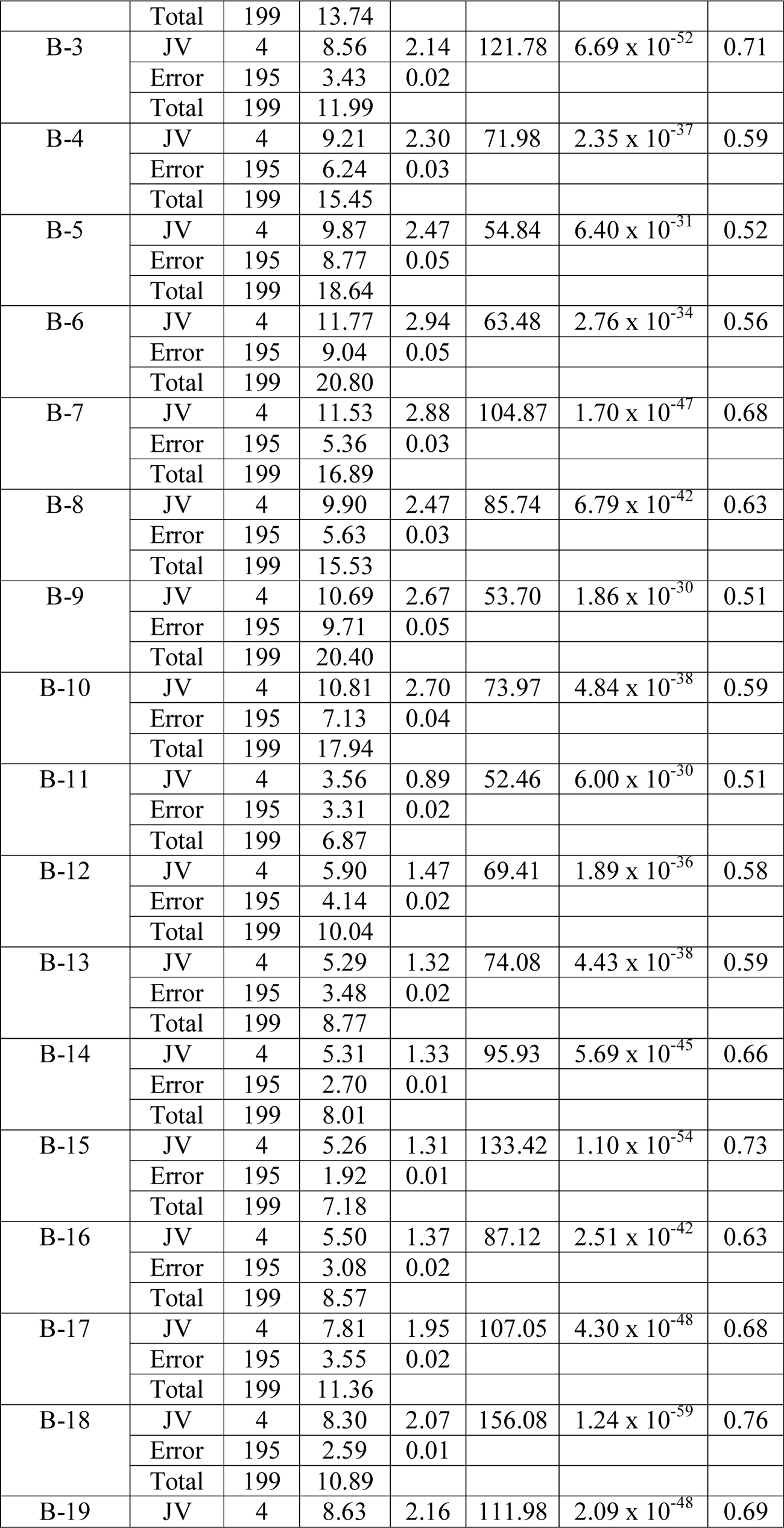
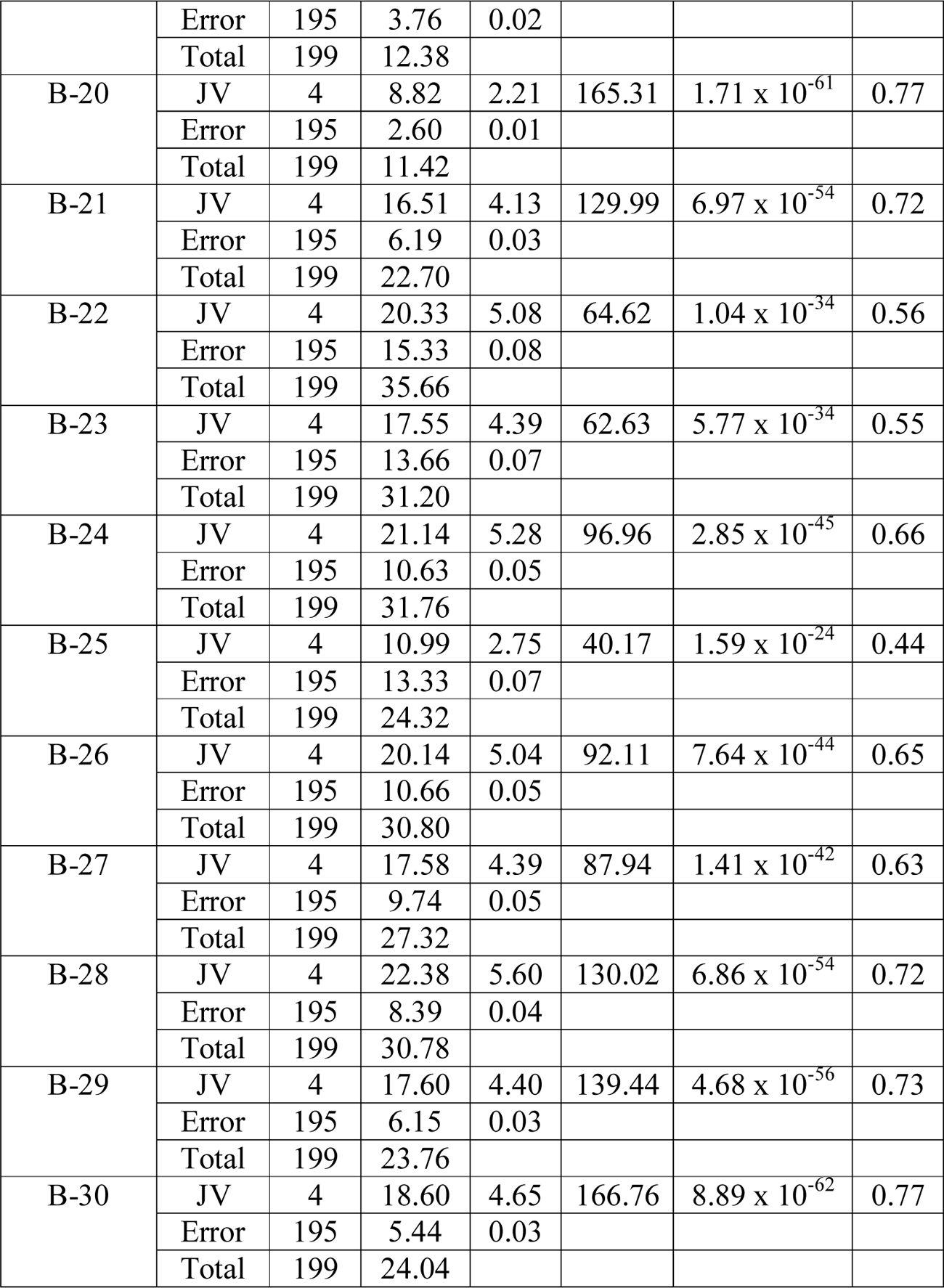
Effect of juice volume on BDM bids in individual sessions. Statistical test: one-way ANOVA. Abbreviations: JV: juice volume, d.f.: degree of freedom, SS: sum of squares, MS: mean square, F: F-statistic, p: p-value, ω^2^: omega-squared effect size.

**Extended Data Table 6-1.**
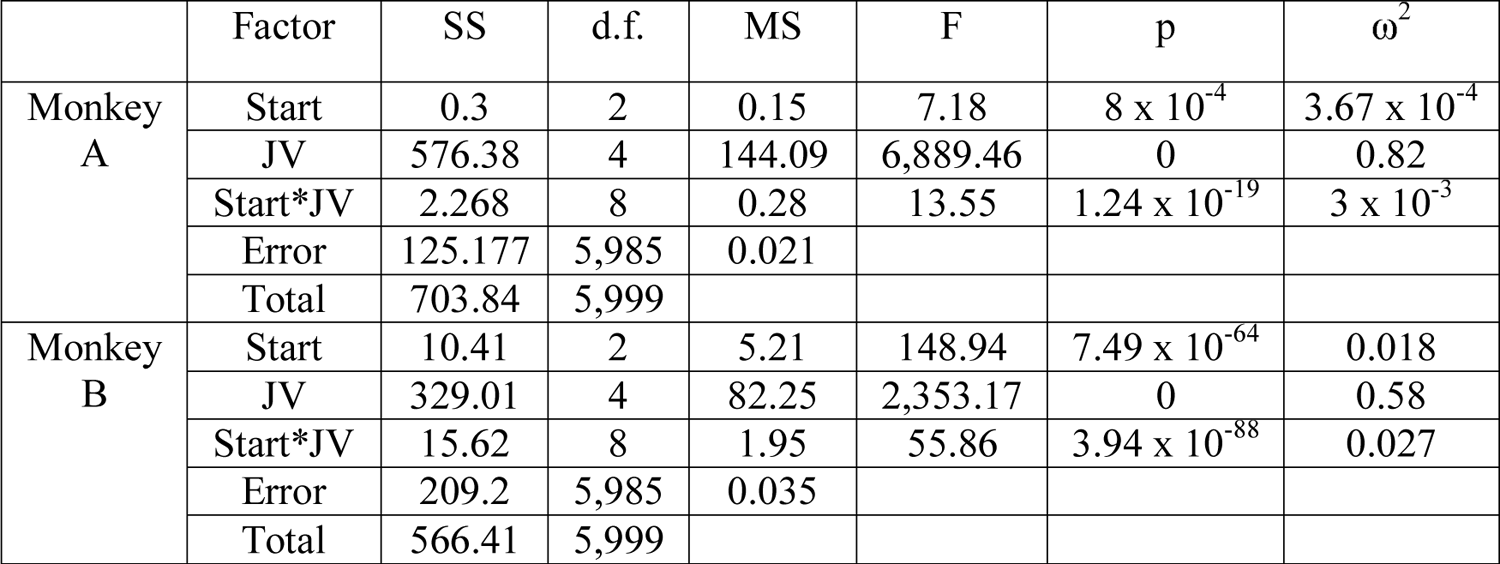
Effects of starting bid position and juice volume on BDM bids. Starting bid position was at bottom, top or random on budget bar. For Monkey A, overall, bids were significantly lower in the top-start BDM than in either the bottom-start (*P* = 6.35 x 10^-4^; unbalanced two-way ANOVA). or random-start versions of the task (*P* = 0.034); for Monkey B, bids were significantly greater in the bottom-start BDM than in either the top-start (*P* = 2.1 x 10^-53^) or random-start versions of the task (*P* = 1.95 x 10^-44^). However, a comparison of effect sizes (ω^2^) reveals that for both monkeys the size of any effect due to starting position, or the interaction of starting position and juice volume, was negligible when compared to that of juice volume alone. Abbreviations: Start: starting bid position, JV: juice bolume, d.f.: degree of freedom, SS: sum of squares, MS: mean square, F: F-statistic, p: p-value, ω^2^: omega-squared effect size.

| | Factor | SS | d.f. | MS | F | p | $\omega^2$ |
| --- | --- | --- | --- | --- | --- | --- | --- |
| Monkey A | Start | 0.3 | 2 | 0.15 | 7.18 | $8 \times 10^{-4}$ | $3.67 \times 10^{-4}$ |
|  | JV | 576.38 | 4 | 144.09 | 6,889.46 | 0 | 0.82 |
| | Start*JV | 2.268 | 8 | 0.28 | 13.55 | $1.24 \times 10^{-19}$ | $3 \times 10^{-3}$ |
|  | Error | 125.177 | 5,985 | 0.021 |  |  |  |
|  | Total | 703.84 | 5,999 |  |  |  |  |
| Monkey B | Start | 10.41 | 2 | 5.21 | 148.94 | $7.49 \times 10^{-64}$ | 0.018 |
|  | JV | 329.01 | 4 | 82.25 | 2,353.17 | 0 | 0.58 |
| | Start*JV | 15.62 | 8 | 1.95 | 55.86 | $3.94 \times 10^{-88}$ | 0.027 |
|  | Error | 209.2 | 5,985 | 0.035 |  |  |  |
|  | Total | 566.41 | 5,999 |  |  |  |  |

**Extended Data Table 6-2.** Alternative models for testing bids as a function of juice volume. While bids capture the subjective value of juice in terms of water given a certain water budget, alone they do not provide a measure of utility. Nevertheless, it is possible that bids are non-linearly related to reward volumes. Here, we compared 4 different models; a simple linear model, a 1-parameter power function, a 2-parameter power function, and the Prelec function (which is usually used for probability distortion but can flexibly fit as either a concave/convex function - similar to power functions - or, as a sigmoid fit). These models were fit to all bids pooled across sessions as mixed effects models with random effects at the session-level. For each model we show the root-mean square error (RMSE); the Akaike information criterion (AIC); the Log likelihood (LL); and Beta coefficients (all p < 0.01). We found the linear model to provide the best fit to the data for both monkeys, as it has the lowest AIC and RMSE. Thus, for further analyses, we performed regressions between bids and reward volumes on individual sessions using linear models. Model data are shown for both monkey A (top) and Monkey B (bottom).

| Monkey A | RMSE | AIC | LL | $\beta_0$ | $\beta_1$ |
| --- | --- | --- | --- | --- | --- |
| **Linear**<br>$Bid = \beta_0 + \beta_1 * RVol$ | 0.14 | -6262.79 | 3136.40 | -0.025 | 1.44 |
| **Simple Power Function**<br>$Bid = RVol^{\beta_0}$ | 0.19 | -2707.02 | 1356.51 | 0.62 | |
| **2-Parameter Power Function**<br>$Bid = \beta_0 * RVol^{\beta_1}$ | 0.14 | -6214.13 | 3112.07 | 1.43 | 1.05 |
| **Prelec Model**<br>$Bid = e^{-\beta_0 * (-\ln(RVol))^{\beta_1}}$ | 0.15 | -5449.00 | 2729.50 | 0.61 | 1.88 |
| Monkey B | RMSE | AIC | LL | $\beta_0$ | $\beta_1$ |
| **Linear**<br>$Bid = \beta_0 + \beta_1 * RVol$ | 0.18 | -3157.18 | 1583.59 | 0.15 | 1.10 |
| **Simple Power Function**<br>$Bid = RVol^{\beta_0}$ | 0.20 | -2110.16 | 1058.08 | 0.55 | |
| **2-Parameter Power Function**<br>$Bid = \beta_0 * RVol^{\beta_1}$ | 0.18 | -3117.69 | 1563.85 | 1.20 | 0.75 |
| **Prelec Model**<br>$Bid = e^{-\beta_0 * (-\ln(RVol))^{\beta_1}}$ | 0.19 | -2789.52 | 1399.76 | 0.54 | 1.37 |

**Extended Data Table 7-1.** BDM bids in common currency of ml of water assessed in the binary choice task. Each table data cell shows ml of water equivalent (mean ± standard deviation) from 200 trials, with number of trails in brackets underneath. B-BDM, T-BDM and R-BDM refer to bid cursor start at bottom, top or random position on the budget bar, respectively.

|  |  | B-BDM | T-BDM | R-BDM | All BDM | All BC |
| --- | --- | --- | --- | --- | --- | --- |
| Monkey A | 0.15 ml | 0.26 ± 0.12<br>(433) | 0.18 ± 0.15<br>(413) | 0.19 ± 0.16<br>(394) | 0.21 ± 0.15<br>(1,240) | 0.25 ± 0.11<br>(400) |
|  | 0.30 ml | 0.37 ± 0.14<br>(400) | 0.36 ± 0.18<br>(376) | 0.35 ± 0.20<br>(392) | 0.36 ± 0.17<br>(1,168) | 0.41 ± 0.16<br>(400) |
|  | 0.45 ml | 0.64 ± 0.16<br>(373) | 0.63 ± 0.14<br>(403) | 0.64 ± 0.18<br>(412) | 0.64 ± 0.16<br>(1,188) | 0.74 ± 0.15<br>(400) |
|  | 0.60 ml | 0.86 ± 0.16<br>(405) | 0.87 ± 0.12<br>(378) | 0.89 ± 0.13<br>(395) | 0.88 ± 0.14<br>(1,178) | 0.98 ± 0.18<br>(400) |
|  | 0.75 ml | 1.02 ± 0.12<br>(389) | 1.03 ± 0.09<br>(430) | 1.07 ± 0.09<br>(407) | 1.04 ± 0.10<br>(1,226) | 1.64 ± 0.34<br>(400) |
| Monkey B | 0.15 ml | 0.40 ± 0.12<br>(398) | 0.35 ± 0.14<br>(406) | 0.21 ± 0.13<br>(422) | 0.32 ± 0.16<br>(1,226) | 0.15 ± 0.10<br>(400) |
|  | 0.30 ml | 0.53 ± 0.18<br>(407) | 0.49 ± 0.14<br>(418) | 0.39 ± 0.24<br>(388) | 0.47 ± 0.20<br>(1,213) | 0.29 ± 0.12<br>(400) |
|  | 0.45 ml | 0.69 ± 0.22<br>(381) | 0.62 ± 0.14<br>(401) | 0.61 ± 0.27<br>(396) | 0.64 ± 0.22<br>(1,178) | 0.52 ± 0.16<br>(400) |
|  | 0.60 ml | 0.86 ± 0.21<br>(417) | 0.73 ± 0.15<br>(379) | 0.84 ± 0.27<br>(390) | 0.81 ± 0.22<br>(1,186) | 0.77 ± 0.18<br>(400) |
|  | 0.75 ml | 1.04 ± 0.16<br>(397) | 0.86 ± 0.12<br>(396) | 1.04 ± 0.20<br>(404) | 0.98 ± 0.18<br>(1,197) | 1.14 ± 0.24<br>(400) |

**Extended Data Table 8-1.** Differences in average per-trial payoff between the simulated optimal bidder and each other bidder. The mean difference in payoff is shown for each reward and each bidder separately. The standard deviation is shown in brackets. Note, data is not shown for the 0.75 ml reward for this animal as the binary choice value for this reward was outside of the bidding range, and so overall payoffs could not be calculated.

|  |  | Reward volume (ml) |  |  |  |
| --- | --- | --- | --- | --- | --- |
|  |  | 0.15 | 0.3 | 0.45 | 0.6 |
| **Bidder** | **Monkey** | 0.0059<br>(0.030) | 0.011<br>(0.036) | 0.028<br>(0.063) | 0.018<br>(0.064) |
|  | **Random** | 0.13<br>(0.18) | 0.085<br>(0.14) | 0.069<br>(0.12) | 0.15<br>(0.20) |
|  | **Over** | 0.024<br>(0.053) | 0.034<br>(0.059) | 0.019<br>(0.044) | 0.0014<br>(0.011) |
|  | **Under** | 0.0024<br>(0.014) | 0.015<br>(0.039) | 0.038<br>(0.061) | 0.02<br>(0.05) |

**Monkey B**
|  |  | Reward volume (ml) |  |  |  |  |
| --- | --- | --- | --- | --- | --- | --- |
|  |  | 0.15 | 0.3 | 0.45 | 0.6 | 0.75 |
| **Bidder** | **Monkey** | 0.028<br>(0.081) | 0.053<br>(0.11) | 0.031<br>(0.07) | 0.018<br>(0.052) | 0.026<br>(0.093) |
|  | **Random** | 0.19<br>(0.23) | 0.13<br>(0.18) | 0.057<br>(0.10) | 0.075<br>(0.13) | 0.22<br>(0.27) |
|  | **Over** | 0.016<br>(0.05) | 0.029<br>(0.057) | 0.034<br>(0.059) | 0.015<br>(0.039) | 0 |
|  | **Under** | 0.00016<br>(0.0022) | 0.0031<br>(0.017) | 0.026<br>(0.051) | 0.035<br>(0.060) | 0.0064<br>(0.031) |

## Notes

### Competing Interest Statement

The authors have declared no competing interest.

### Summary of Updates

Revision in wording and figures.

## References

1. Bateman IJ, Munro A, Rhodes B, Starmer C, Sugden R (1997) Does Part-Whole Bias Exist? An Experimental Investigation. The Economic Journal 107:322–332.

2. Becker GM, DeGroot MH, Marschak J (1964) Measuring utility by a single response sequential method. Behav Sci 9:226–232.

3. Brainard DH (1997) The psychophysics toolbox. Spatial Vision 10:433–436.

4. Brandstätter E, Gigerenzer G, Hertwig R (2006) The priority heuristic: Making choices without trade-offs. Psychological Review 113:409–432.

5. Bujold PM, Ferrari-Toniolo S, Chi U, Seak L, Schultz W (2021) Comparing utility functions between risky and riskless choice in rhesus monkeys. Anim Cog (in press).

6. Chib VS, Rangel A, Shimojo S, O’Doherty JP (2009) Evidence for a common representation of decision values for dissimilar goods in human ventromedial prefrontal cortex. J Neurosci 29:12315–12320.

7. Genest W, Stauffer WR, Schultz W (2016) Utility functions predict variance and skewness risk preferences in monkeys. Proc Natl Acad Sci (USA) 113:8402–8407.

8. Harris A, Adolphs R, Camerer C, Rangel A (2011) Dynamic construction of stimulus values in the ventromedial prefrontal cortex. PLoS One 6:e21074.

9. Hayden BY, Heilbronner SR, Platt ML (2010) Ambiguity aversion in rhesus macaques. Front Neurosci 4:166.

10. Karni E, Safra Z (1987) Preference reversals and the observability of preferences by experimental methods. Econometrica 55:675–685.

11. Knetsch JL, Sinden JA (1984) Willingness to Pay and Compensation Demanded: Experimental Evidence of an Unexpected Disparity in Measures of Value. Quart J Econ 99:507–521.

12. Kobayashi S, Carvalho OP, Schultz W (2010) Adaptation of Reward Sensitivity in Orbitofrontal Neurons. J Neurosci 30:534–544.

13. Kobayashi S, Schultz W (2008) Influence of reward delays on responses of dopamine neurons. J Neurosci 28:7837–7846.

14. Lak A, Stauffer WR, Schultz W (2014) Dopamine prediction error responses integrate subjective value from different reward dimensions. Proc Natl Acad Sci (USA) 111:2343–2348.

15. Linder NS, Uhl G, Fliessbach K, Trautner P, Elger CE, Weber B (2010) Organic labeling influences food valuation and choice. NeuroImage 53:215–220.

16. Louie K, Grattan LE, Glimcher PW (2011) Reward Value-Based Gain Control: Divisive Normalization in Parietal Cortex. J Neurosci 31:10627–10639.

17. Lusk JL, Alexander C, Rousu MC (2007) Designing Experimental Auctions for Marketing Research: The effect of Values, Distributions, and Mechanisms on Incentives for Truthful Bidding. Rev Mark Sci 5:1.

18. Lusk JL, Shogren JF (2007) Experimental auctions: Methods and applications in economic and marketing research. Cambridge University Press.

19. Milgrom PR, Weber RJ (1982) A theory of auctions and competitive bidding. Econometrica 50:1089–1122.

20. Moldovanu B, Tietzel M (1998) Goethe’s Second-Price Auction. J Polit Econ 106:854–859.

21. Montague PR, Berns GS (2002) Neural economics and the biological substrates of valuation. Neuron 36:265-284.

22. Padoa-Schioppa C, Assad JA (2006) Neurons in the orbitofrontal cortex encode economic value. Nature 441:223–226.

23. Padoa-Schioppa C, Rustichini A (2014) Rational Attention and Adaptive Coding: A Puzzle and a Solution. Am Econ Rev 104:507–513.

24. Pastor-Bernier A, Stasiak A, Schultz W (2019) Orbitofrontal signals for two-component choice options comply with indifference curves of Revealed Preference Theory. Nat Comm 10:4885.

25. Piantadosi ST, Hayden BY (2015) Utility-free heuristic models of two-option choice can mimic predictions of utility-stage models under many conditions. Front Neurosci 9:105.

26. Plassmann H, O’Doherty J, Rangel A (2007) Orbitofrontal cortex encodes willingness to pay in everyday economic transactions. J Neurosci 27:9984–9988.

27. Platt ML, Glimcher PW (1999) Neural correlates of decision variables in parietal cortex. Nature 400:233–238.

28. Platt ML, Huettel SA (2008) Risky business: the neuroeconomics of uncertainty. Nat Neurosci 11:398–403.

29. Prelec D (1998) The Probability Weighting Function. Econometrica 66:497–527.

30. Rieskamp J, Busemeyer JR, Mellers B (2006) Extending the Bounds of Rationality: Evidence and Theories of Preferential Choice. J Econ Lit 44:631–661.

31. Samejima K, Ueda Y, Doya K, Kimura M (2005) Representation of action-specific reward values in the striatum. Science 310:1337–1340.

32. Seo H, Lee D (2009) Behavioral and neural changes after gains and losses of conditioned reinforcers. J Neurosci 29:3627–3641.

33. Soltani A, De Martino B, Camerer C (2012) A Range-Normalization Model of Context-Dependent Choice: A New Model and Evidence. PLoS Comput Biol 8:e1002607.

34. Stauffer WR, Lak A, Schultz W (2014) Dopamine reward prediction error responses reflect marginal utility. Curr Biol 24:2491–2500.

35. Tang DW, Fellows LK, Dagher A (2014) Behavioral and neural valuation of foods is driven by implicit knowledge of caloric content. Psychol Sci 25:2168–2176.

36. Tobler PN, Fiorillo CD, Schultz W (2005) Adaptive Coding of Reward Value by Dopamine Neurons. Science 307:1642–1645.

37. Tversky A (1972) Elimination by aspects: A theory of choice. Psychological Review 79:281–299.

38. Tversky A, Simonson I (1993) Context-Dependent Preferences. Managemt Sci 39:1179-1189.

39. Tymula A, Woelbert E, Glimcher P (2016) Flexible valuations for consumer goods as measured by the Becker-DeGroot-Marschak mechanims. J Neurosci Psychol Econ 2:65–77.

40. Tyson-Carr J, Kokmotou K, Soto V, Cook S, Fallon N, Giesbrecht T, Stancak A (2018) Neural correlates of economic value and valuation context. J Neurophysiol 119:1924–1933.

41. Vickrey W (1961) Counterspeculation, Auctions, and Competitive Sealed Tenders. J Finance 16:8–37.

42. Vlaev I, Chater N, Stewart N, Brown GDA (2011) Does the brain calculate value? Trends Cog Sci 15:546–554.

